# The Hitchhiker’s Guide to Sequencing Data Types and Volumes for Population-Scale Pangenome Construction

**DOI:** 10.1101/2024.03.14.585029

**Authors:** Prasad Sarashetti, Josipa Lipovac, Filip Tomas, Mile Šikic, Jianjun Liu

**Affiliations:** Laboratory of Human Genomics, Genome Institute of Singapore, A*STAR, Singapore, Singapore; Laboratory for Bioinformatics and Computational Biology, Faculty of Electrical Engineering and Computing, University of Zagreb, Zagreb, Croatia; Laboratory of AI in Genomics, Genome Institute of Singapore, A*STAR, Singapore, Singapore; Yong Loo Lin School of Medicine, National University of Singapore, Singapore

**Keywords:** LRS Special Issue, Pangenome, De Novo Assembly, Sequencing Platforms, Population-Level Studies

## Abstract

Long-read (LR) technologies from Pacific Biosciences (PacBio) and Oxford Nanopore Technologies (ONT) have transformed genomics research by providing diverse data types like HiFi, Duplex, and ultra-long ONT (ULONT). Despite recent strides in achieving haplotype-phased gapless genome assemblies using long-read technologies, concerns persist regarding the representation of genetic diversity, prompting the development of pangenome references. However, pangenome studies face challenges related to data types, volumes, and cost considerations for each assembled genome, while striving to maintain sensitivity. The absence of comprehensive guidance on optimal data selection exacerbates these challenges. To fill this gap, our study evaluates available data types, their significance, and the required volumes for robust de novo assembly in population-level pangenome projects. The results show that achieving chromosome-level haplotype-resolved assembly requires 20x high-quality long reads (HQLR) such as PacBio HiFi or ONT duplex, combined with 15-20x of ULONT per haplotype and 30x of long-range data such as Omni-C. High-quality long reads from both platforms yield assemblies with comparable contiguity, with HiFi excelling in NG50 and phasing accuracies, while usage of duplex generates more T2T contigs. As Long-Read Technologies advance, our study reevaluates recommended data types and volumes, providing practical guidelines for selecting sequencing platforms and coverage. These insights aim to be vital to the pangenome research community, contributing to their efforts and pushing genomic studies with broader impacts.

## Introduction

A high-quality and complete human reference genome is the fundamental bedrock supporting genetic studies of human diseases and population structures. Over the past two decades, the human reference genome employed in genetic studies has been meticulously crafted from genomic segments sourced from thousands of individuals (Lander et al. 2001; Schneider et al. 2017). Despite efforts to assemble high-quality, gapless genomes such as T2T-CHM13 (Nurk et al. 2022), T2T-YAO (He et al. 2023), CN1 (Yang et al. 2023), I002C (https://github.com/LHG-GG/I002C) or HG002 (https://github.com/marbl/HG002), such references raise concerns regarding their abilities to represent genetic variations across diverse human populations accurately. The prevailing consensus is that no singular reference sequence can adequately encapsulate the complex genomic diversity across global populations (Yang et al. 2019). This understanding highlights the crucial need for high-quality reference genome panels that accurately resolve haplotypes, presenting the complex genetic variations observed within distinct populations (Lou et al. 2022; Deng et al. 2022; Ballouz et al. 2019). In parallel, there is a growing trend to shift from a singular reference to a pangenomic approach, which supports a broader range of genomic diversity, acknowledging the complexities within and across diverse human populations (Sherman and Salzberg 2020; Wang et al. 2022; Liao et al. 2023; Gao et al. 2023). This shift is supported by the rapid advancement of computational methodologies for de novo genome assembling analysis (Garrison et al. 2018; Eizenga et al. 2020; Li et al. 2020; Vernikos 2020; Ebler et al. 2022; Hickey et al. 2023).

Haplotype-resolved genome sequences are the building blocks for pangenome construction. However, despite the contradictory nature of cost and sensitivity, both of which play vital roles in pangenomic projects, most recent studies (Liao et al. 2023; Gao et al. 2023) lack a comprehensive evaluation and guidelines related to the optimal data types and volumes required, mostly relying on the propositions of assembly tool authors for volume and data type requirements.

At the forefront of long-read technology (LRT) innovation, Pacific Biosciences (PacBio) and Oxford Nanopore Technologies (ONT) stand out as the primary driving forces, spearheading advancements in this field through their groundbreaking contributions. PacBio’s long reads (LR) have excelled in read quality, while ONT has leveraged its competitive edge in providing substantial read lengths at a lower cost (Logsdon et al. 2020). To address the disparity between read quality and length, ONT has recently introduced a novel technique termed “Duplex”, capable of achieving a quality level of Q30, thereby bridging the gap between read quality and read length (https://nanoporetech.com/about-us/news/oxford-nanopore-tech-update-new-duplex-method-q30-nanopore-single-molecule-reads-0). Recent comparisons indicate similar performance between the two platforms in structural variation (SV) analysis (Du et al. 2022; Harvey et al. 2023) (details of difference studies utilizing longs are discussed elsewhere (Coster et al. 2021)), but the performance for de novo genome assembling analysis has not been properly evaluated and compared.

In this study, we evaluated different data types and the minimum data volume required to establish a robust pipeline of de novo assembly for population-level pangenome projects. Specifically, we conducted a performance comparison between ONT’s Duplex dataset and PacBio HiFi dataset. In this comparison, we extensively examined the performance of these datasets in the context of de novo genome assembly, scrutinizing their effectiveness in achieving high-quality phased genomes with enhanced contiguity and completeness. Given the swift advancements in Long-Read Technologies (LRT), it’s prudent to reassess the recommended data types and volume outlined in this study, aligning them with the practical economic limitations within the scope of your research endeavours.

## Results

### DNA Sequencing

The data used for this research are generated as a part of an ongoing effort to generate telomere-2-telomere diploid assembly of a male Singaporean of Indian ancestry I002C(https://github.com/LHG-GG/I002C/ https://github.com/lbcb-sci/I002C). Through sequencing on various platforms, we obtained the following dataset for the child sample: 152.97 Gb (∼50.99x) PacBio HiFi data, 193.66 Gb (∼64.55x) ONT Duplex data, 441.07 Gb (∼147x) ONT ultralong data (ULONT) and 222.21 (∼74.07x) Omni-C data. For the paternal sample-107.69 Gb (∼35.90x) and maternal sample-112.48 Gb (∼37.49x) MGI paired-end data was sequenced [Table 1]. On average, the Duplex reads were twice as long as the HiFi reads, yet they maintained a comparable level of read quality [Figure 1].

**Figure 1.**
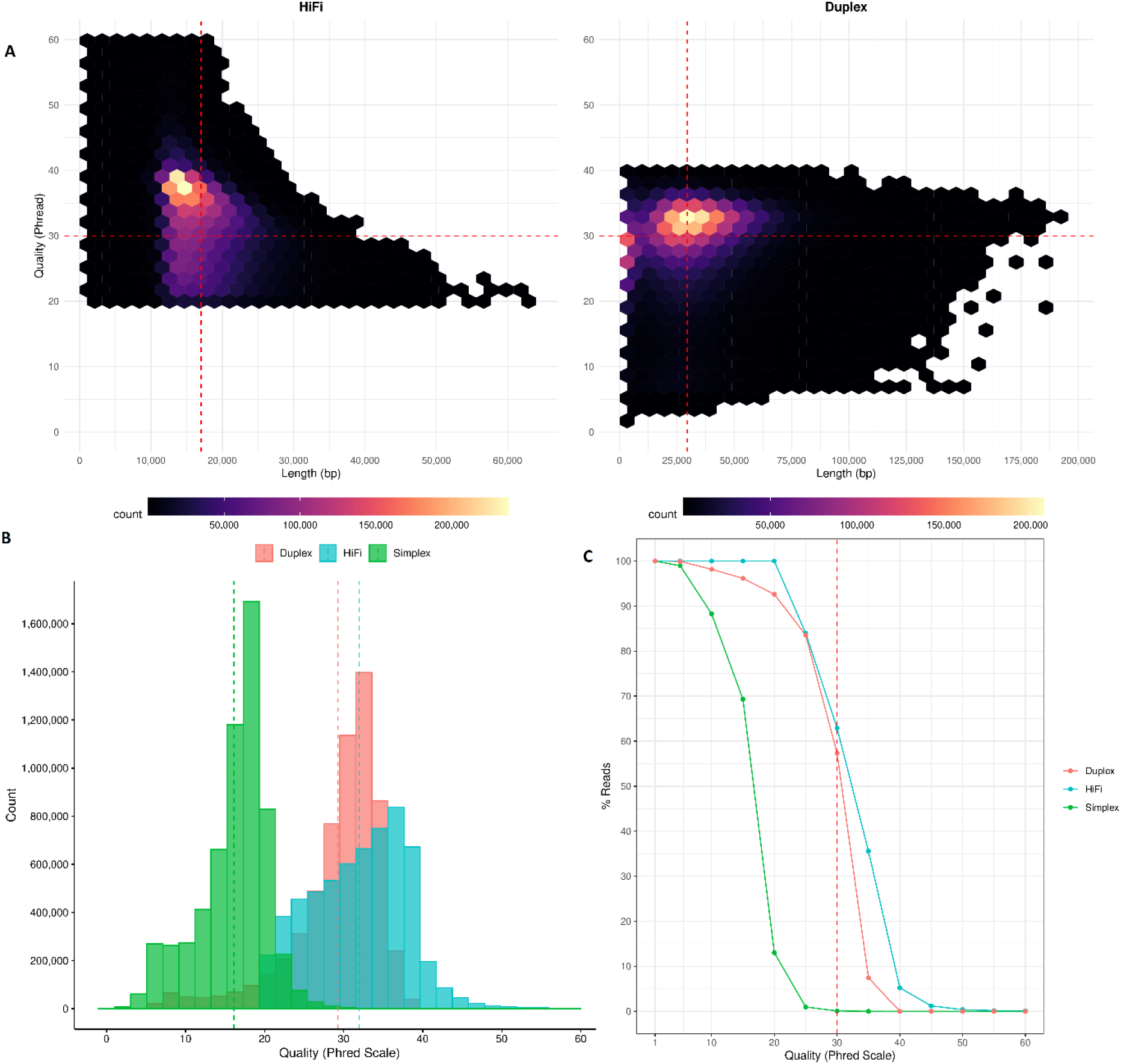
Comparison of read length and quality (Phred scale) between PacBio HiFi and ONT Duplex **A)** Distribution of read length and quality of HiFi and Duplex reads. **B)** Comparison of read quality among ONT Simplex Vs ONT Duplex Vs PacBio HiFi **C)** Percentage of reads which are Q30 and above.

**Table 1:** Summary of sequencing data from different platforms.

| Sample | Platform | Data type | Reads | Total bases (Gb) | Depth | Min Length | Max Length | Average Read Length | N50 | %GC |
| --- | --- | --- | --- | --- | --- | --- | --- | --- | --- | --- |
| Child | PacBio | HiFi | 8,986,857 | 152,977,790,522 | 50.99 | 8 | 63,921 | 17,022.40 | 17,132 | 40.78 |
|  | ONT | Duplex | 6,561,950 | 193,663,321,924 | 64.55 | 1 | 191,644 | 29,513.10 | 38,128 | 40.85 |
|  | ONT | Ultralong | 12,710,241 | 441,077,651,554 | 147.03 | 1 | 1,434,925 | 34,702.50 | 87,761 | 40.58 |
|  | MGI | Omni-C | 1,476,892,046 | 222,216,283,752 | 74.07 | 150 | 150 | 150 | 150 | 42 |
| Paternal | MGI | Short reads | 717,979,504 | 107,696,925,600 | 35.9 | 150 | 150 | 150 | 150 | 40.48 |
| Maternal | MGI | Short reads | 749,860,990 | 112,479,148,500 | 37.49 | 150 | 150 | 150 | 150 | 40.28 |

Note: The coverage depth calculation was based on a genome size estimation of 3 Gb. PacBio-
Pacific Biosciences; HiFi- High Fidelity reads; ONT- Oxford Nanopore Technologies; Gb- Gigabases

### Coverage saturation analysis of population-scale de novo assembly

To leverage the potential of long reads, the assembling analysis specifically utilized high-quality long reads (HQLR) such as HiFi and Duplex, which are 10 kb or longer and ULONT reads of at least 100 kb [Table S2]. We examined the importance of diverse data types and offered general observations on the sequencing depth or data volume required for de novo assembling analyses at scale.

#### 1) Data down-sampling

We generated down-sampled data sets for different data types at specific depths considering a haploid genome size of 3 Gbp [Table S3-S5].

i. HiFi and Duplex reads -- 20, 30, 35, 40, 45, and 50
ii. ULONT – 10, 20, 30, 40, 50, and 60
iii. Omni-C – 10, 20, and 30

Owing to the longer length of Duplex reads, achieving the same sequencing depth requires, on average, twice as many HiFi reads as Duplex reads at any given coverage level, as demonstrated in [Figure S1].

#### 2) Evaluation of assembly results in terms of sequence saturation

To evaluate the impact of sequencing coverage on assembling performance and identify the coverage saturation point where assembly contiguity begins to plateau, we utilized Hifiasm to assemble HiFi/Duplex data independently (HQLR_Only) and in conjunction with ULONT data across varying coverage depths. The assembly results show a clear positive correlation between the augmentation of data coverage (HiFi/Duplex) and assembly performance for both primary assemblies [Figure 2], representing a mosaic of the two haplotypes and the two haplotypes derived from the phased assembly.

**Figure 2.**
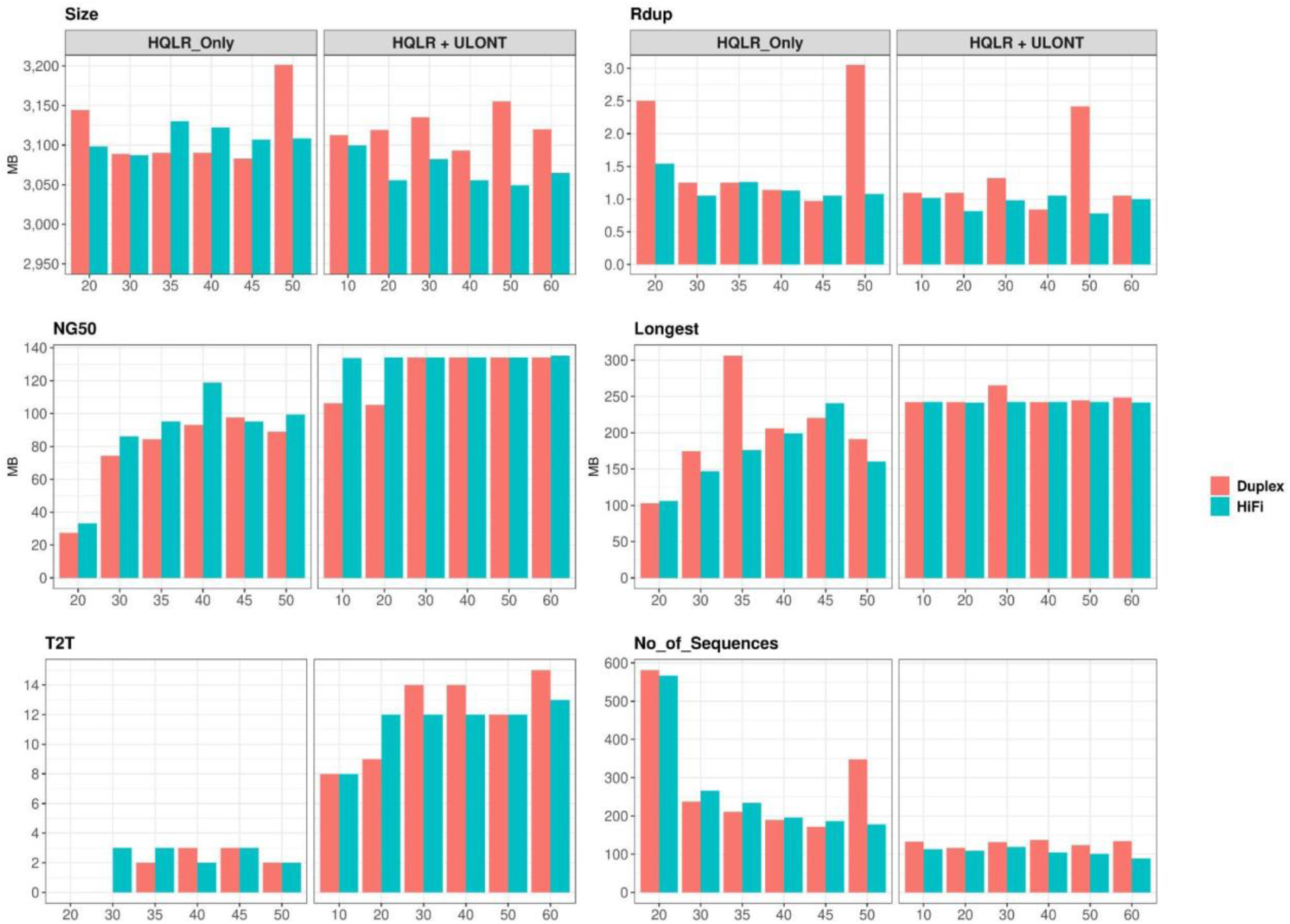
Comparison of assembly performance versus data coverage for the primary assembly. “HQLR_Only” denotes assemblies generated solely with HiFi or Duplex data across various coverages. “HQLR_ULONT” signifies assemblies generated with a saturation coverage (35x) of HiFi and Duplex data combined with various ULONT coverages.

At any given coverage, the assembled genome size aligns well with expected genome sizes (2.9Gb paternal, 3Gb maternal, and 3.1Gb primary assembly). The inflated assembled genome size positively correlated with the duplication rate (Rdup). As the data coverage increases, key assembly contiguity features such as NG50, Longest contig length, and Telomere-2-Telomere [T2T] contigs exhibit an upward trend, while the “No_of_Sequences” demonstrate a downward trajectory. Assembly contiguity reaches plateaus when the HQLR-only (HiFi/Duplex) coverage exceeds 35x.

Furthermore, in combination with ULONT data, even as low as 10x ULONT along with 35x of HQLR plateau coverage significantly enhances assembly contiguity compared to that of 50x HQLR-only assembly. The inclusion of ULONT data notably improves the assembly of telomere-to-telomere contigs. Assembly contiguity reaches a plateau with ULONT coverage exceeding 30x. We observed a similar trend for two haplotype-resolved assemblies [Figure S2]. The detailed assembly statistics are provided in the Supplementary Materials [Table S6-S11]. Hereafter, in the figures and tables, “ HQLR + ULONT” denotes HQLR (HiFi/Duplex) coverage of 35x, representing the HQLR-only plateau coverage, combined with various ULONT coverages. Similarly, “ HQLR + ULONT + Omni-C” signifies 35x HQLR coverage combined with 30x ULONT coverage, representing the ULONT plateau coverage, along with different levels of Omni-C coverage.

#### 3) Improvement of phasing with Omni-C

The Hifiasm tool is capable of producing pseudo-haplotypes or a dual assembly using HiFi/Duplex data alone or in conjunction with ULONT. This process efficiently captures the heterozygous variances across the two haplotypes. HiFi only assemblies demonstrate relatively fewer switch errors owing to their higher quality compared to Duplex only assemblies. Conversely, the longer read lengths of Duplex data contribute to achieving superior global phasing (hamming) compared to HiFi reads [Figure S3]. Like Duplex, but even more significantly, due to their lengths, ULONT reads improve phasing (Jain et al. 2018). However, even with ULONT reads of significant length, assemblers generate contigs with short phase blocks that often show increased phasing errors [Figure 3].

**Figure 3.**
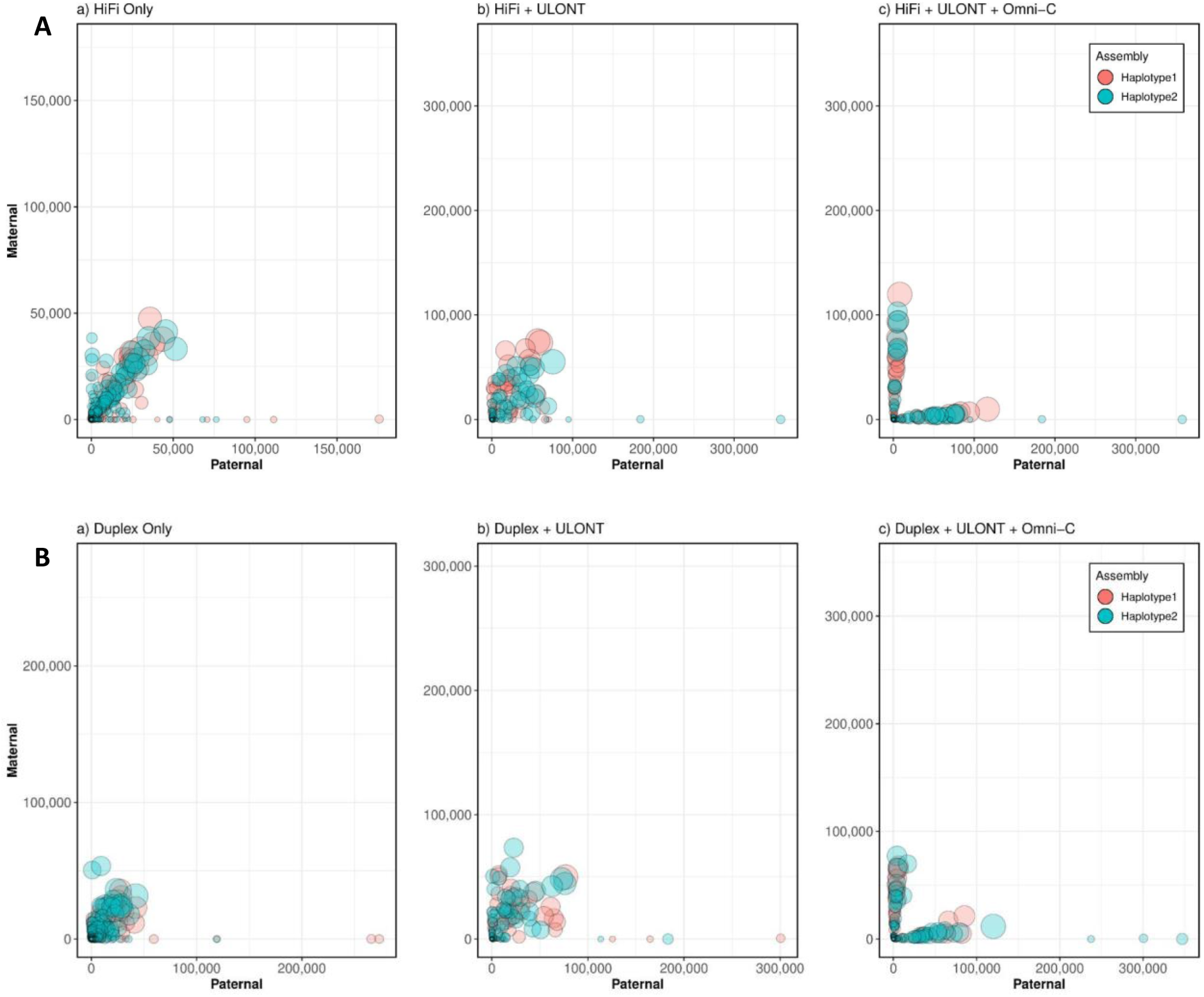
Comparison of phasing accuracies of different assemblies (A - HiFi assembly, B - Duplex assembly). a) Phasing accuracy of dual assembly generated from data-only (HiFi/Duplex) b) Phasing accuracy of dual assembly in conjunction with ULONT c) Haplotype separated assemblies with Omni-C data. Each circle denotes a contig, size reflecting its length. Circle’s positions are determined by the number of maternal and paternal k-mers derived from high-quality short reads on respective contigs. Contigs positioned along the axis indicate higher phasing accuracy.

Incorporating even low coverage of long-range chromatin interaction data like Omni-C, such as 10x, results in a notable reduction in globally incorrectly phased variants (measured by hamming error), leveraging the long-range chromatin interaction information provided by Omni-C. Even though long-range interaction data can produce full-length phased contigs from different chromosomes, contigs from maternal and paternal origin can be mixed in one haplotype [Figure 3]. This intrinsic ambiguity in long-range interaction data phasing is attributed to the challenge of identifying markers that define the paternal and maternal origin, a task not easily achievable with offspring data alone, except in the case of sex chromosomes. Despite this improvement, switch errors, which measure the local inaccuracies of heterozygous variants, remain largely unaffected due to the limitations in the information offered by Omni-C reads. Besides its phasing capability, Omni-C data can also be utilized for scaffolding. Omni-C coverage saturation concerning phasing can be observed at 10x. Since Hifiasm does not leverage long-range data for scaffolding to enhance contiguity, higher coverage may prove advantageous for scaffolding processes. Determining the optimal coverage for long-range chromatin interaction data (Omni-C/Hi-C) is beyond the scope of this study, as discussed elsewhere (Sur et al. 2022).

#### 4) Genome Completeness and Quality

Genome completeness assessed through single copy gene analysis revealed that assemblies from data-only exhibited slightly lower performance with an average of 98.32% single copy, 1.46% duplicated, 0.18% fragmented, and 0.03% missing genes [Figure 4a]. Meanwhile, assemblies generated with HQLR + ULONT data showed higher completeness values with 98.54% single copy, 1.26% duplicated, 0.17% fragmented, and 0.03 missing genes [Figure 4b]. Increased coverage resulted in marginal improvements in gene completeness, with HQLR-only assemblies reaching saturation around 35x coverage, and ULONT assemblies around 30-40x [Table S12]. Results for haplotype-resolved assemblies don’t follow a clear trend but show noticeable improvements with the incorporation of ULONT reads [Figure S4-S5, Table S13-S14].

**Figure 4.**
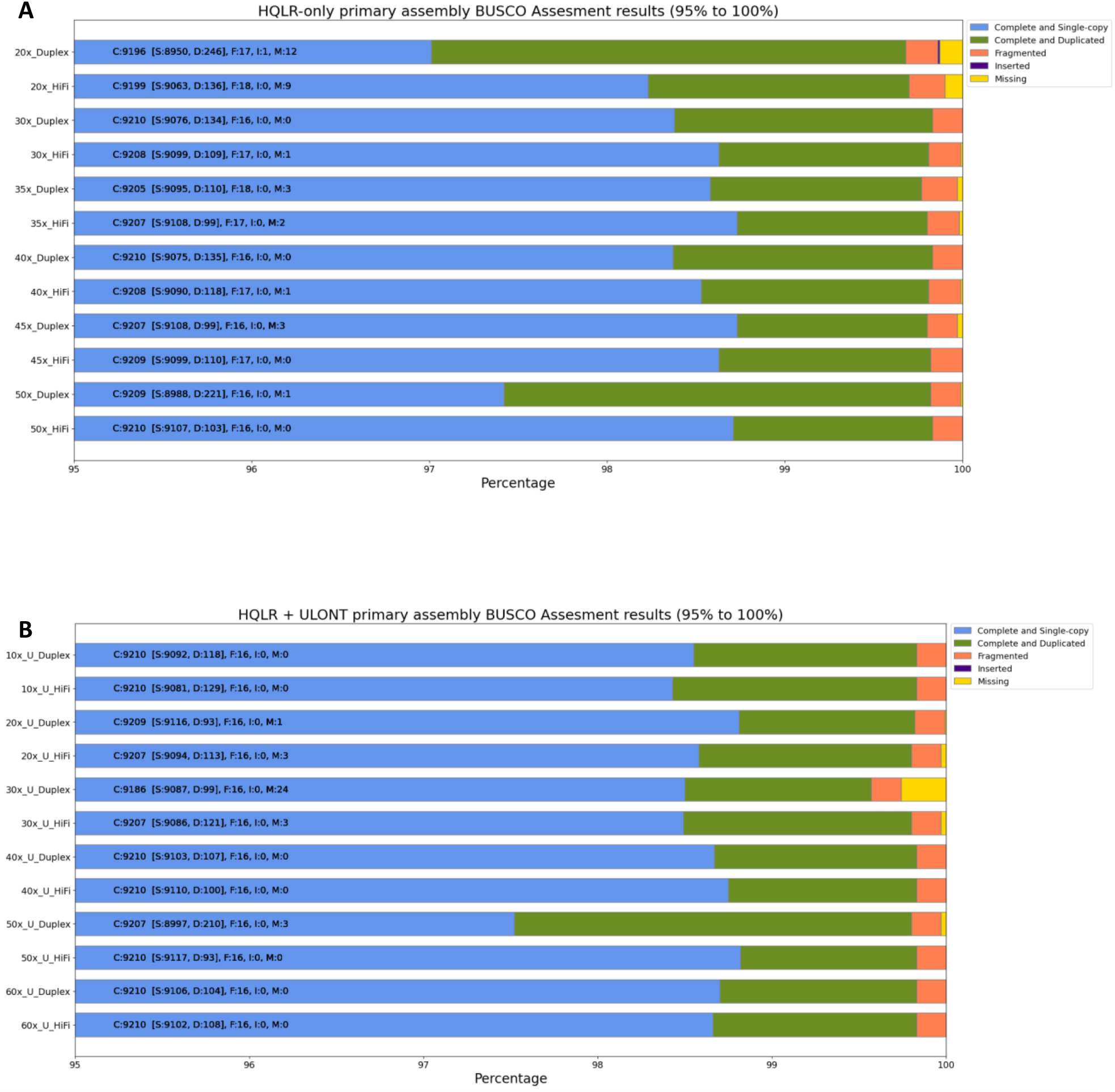
Assessment of gene completeness analysis. A) Output from the assemblies generated from HQLR-only B) Output from assemblies generated from HQLR + ULONT

The estimated k-mer completeness, which indicates the proportion of reliable k-mers from the reads found in the assembly, averaged 96% for the primary assemblies. This finding aligns with the gene completeness analysis results presented in [Figure 5]. Haplotype-resolved assemblies showed a marginally lower average of 95% [Table S15, Figure S6]. The genome completeness analysis showed slightly better results for assemblies from HiFi data compared to Duplex data, both in single-copy analysis and k-mer-based analysis.

**Figure 5.**
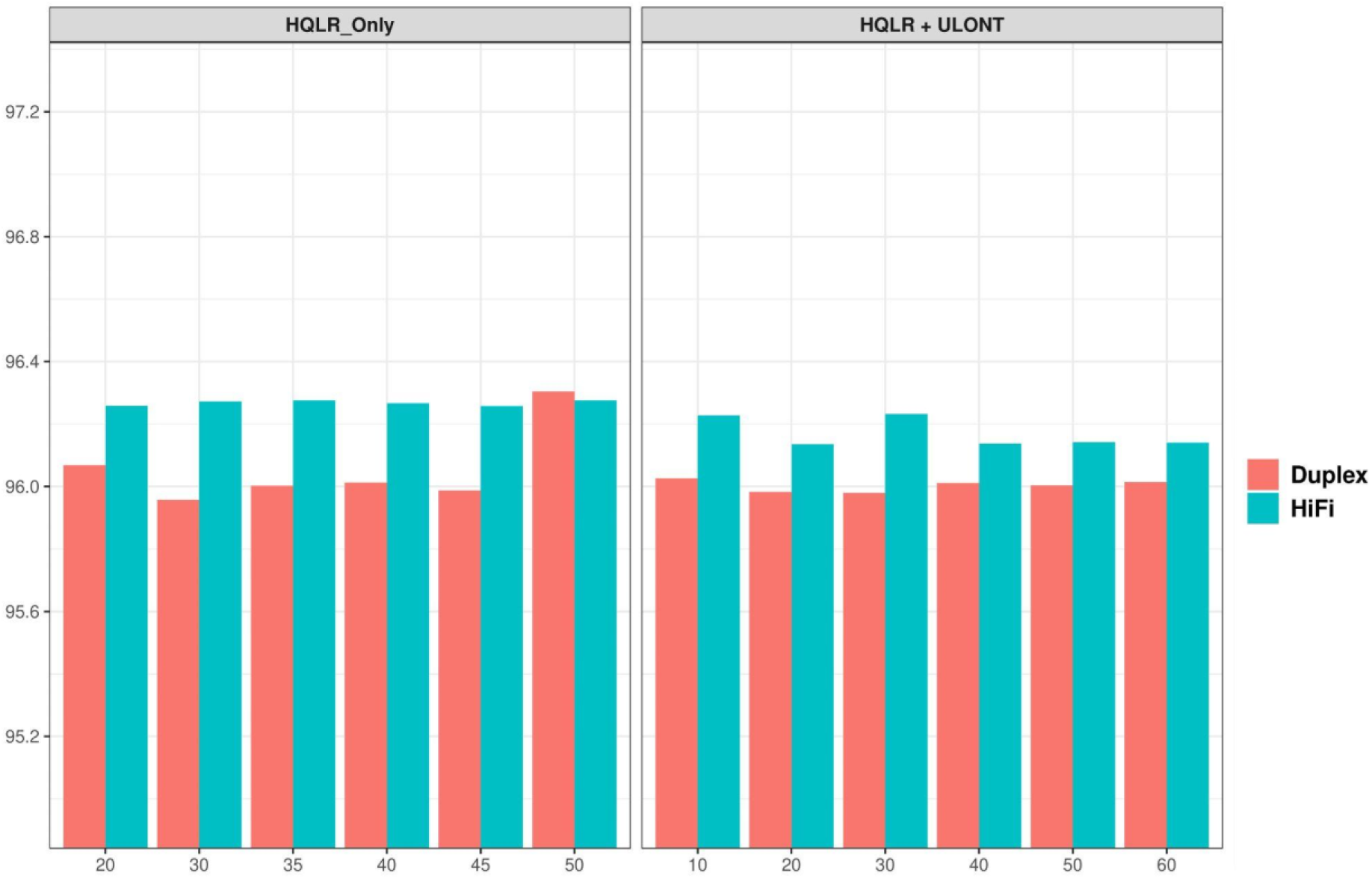
K-mer-based genome completeness analysis.

Assembly quality assessed from k-mers as measured by phred scale quality score (QV) generally showed improvement with increased coverage for both primary assemblies [Figure 6, Table S16] and haplotype-resolved assemblies [Figure S7, Table S16].

**Figure 6.**
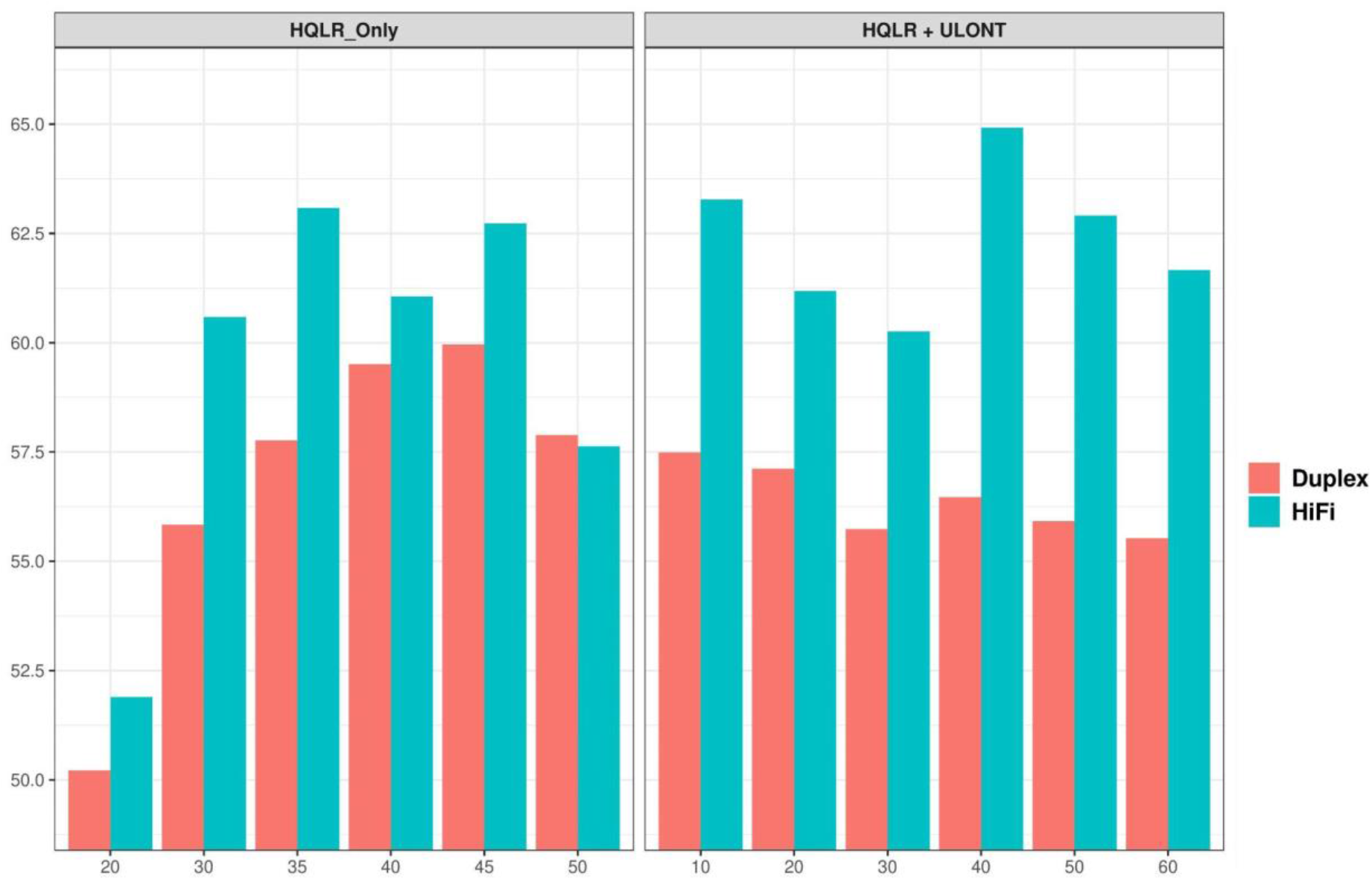
K-mer-based genome quality scores.

#### 5) Computational requirements

We conducted comparisons of both the runtime and peak memory consumption of assembling steps across various coverage levels for specific data types and assemblies resulted from the various combinations of different data types. The computational demands are of paramount importance, particularly in studies conducted at population scale and those utilizing cloud-based platforms for analysis.

The error correction process is a critical and most time-intensive step taking more than half of the total execution time, followed by the graph construction by long read assemblers. By default, Hifiasm performs three rounds of error correction of input HiFi/Duplex reads. Consequently, the time and memory requirements exhibit an upward trajectory with increasing coverage when assemblies are derived solely from data. In the case of “HQLR + ULONT”, where fixed data is employed, in this study 35x of HiFi/Duplex represents a plateau coverage, computational time shows an upward trend with the increased coverage of ULONT, while memory requirements remain stable across coverage levels. This stability in memory consumption is attributed to the implementation of ULONT data in their algorithm (Cheng et al. 2023).

The incorporation of long-range chromatin interaction data (Omni-C) primarily utilized for phasing and resolving graph tangles reveals that both time and memory requirements remain more consistent across increased long-range data coverage [Figure 7, Table S17].

**Figure 7.**
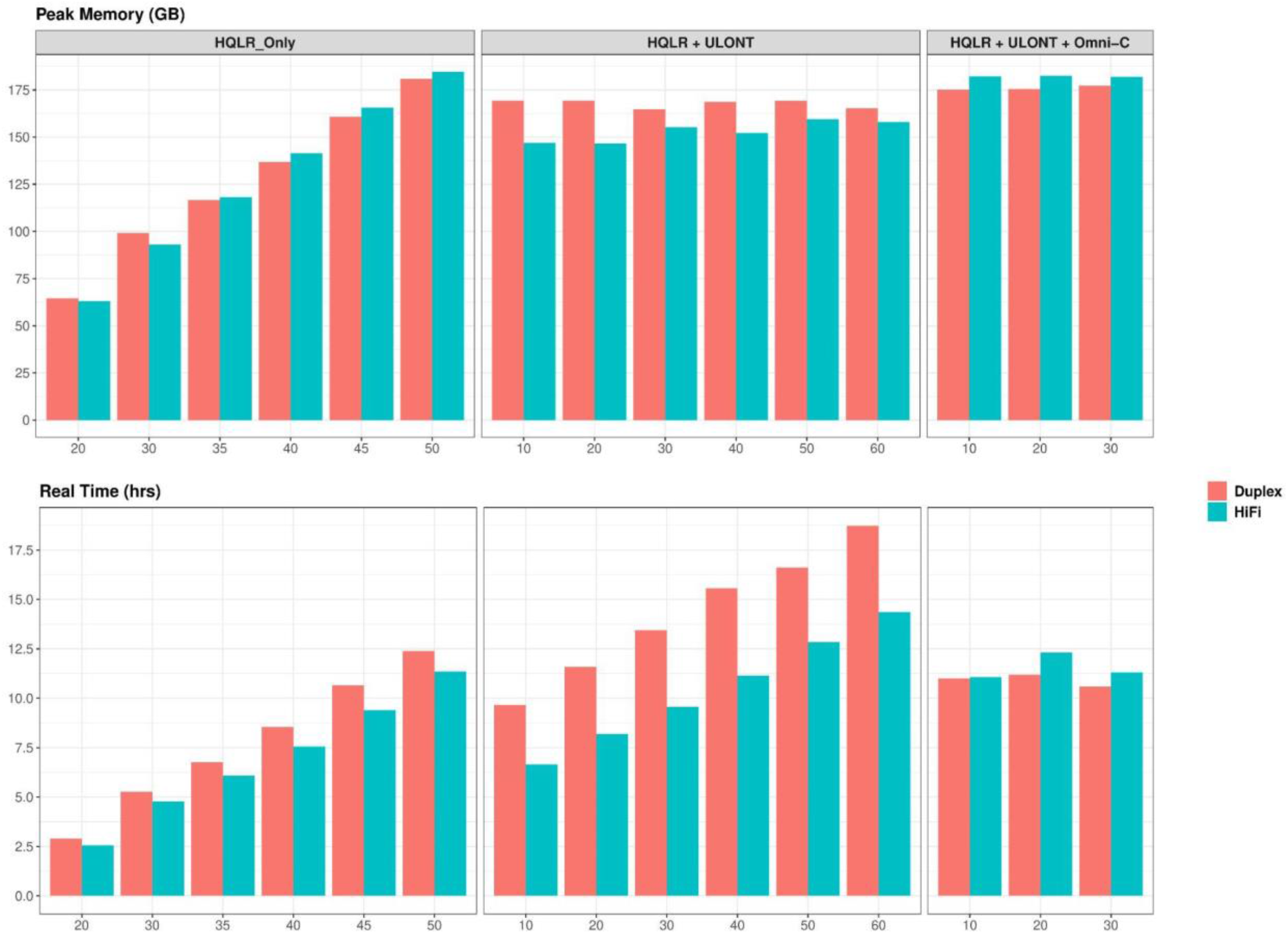
Computation resources consumed by Hifiasm across different data types and coverages

### Comparison of HiFi and Duplex reads performance in *de novo* assembly

As pioneers in long-read technology (LRT), PacBio and ONT continually refine their technologies and develop new advancements to deliver high-quality data at increasingly affordable prices. The recent launch of the PacBio Revio platform (https://www.pacb.com/revio/) stands as a testament to this commitment, elevating HiFi yield by 15x while maintaining impeccable data quality compared to its predecessor, the PacBio Sequel II platform. The assembly contiguity achieved with HiFi data exhibits nearly identical performance on both the Sequel IIe and Revio platforms (Harvey et al. 2023). This substantial boost in data yield has effectively mitigated affordability concerns in comparison to competition. Similarly, ONT has unveiled the enhanced R10 flowcell and introduced the innovative “Duplex” method, which achieves read quality nearing Q30 by sequencing both the template and complement strands of a single molecule. The effectiveness of these cutting-edge data types has been demonstrated in variant calling (Harvey et al. 2023) and methylation studies (Ni et al. 2023), showcasing their utility and performance across different genomic applications.

Here, we conducted a comparison between the new data types regarding their performance in de novo assembly using the current and publicly available HG002 dataset(45x Duplex, 67x HiFi, 30x of ULONT, HiC). HG002 Duplex and HiFi data from the Revio platform were downloaded from HPRC (https://humanpangenome.org/data). The assemblies were constructed with the same coverage (i.e. 35x HiFi/Duplex + 30x ULONT + 30x Hi-C [HG002]/Omni-C [I002C]) data with default parameters of Hifiasm across three independent replicates. We assigned a rank of 1 to the highest value and 0 otherwise for each assembly feature. The sum of these ranks was then computed for both HiFi and duplex assemblies to evaluate their performance based on specific criteria, ranging from best to worst [Figure 8]. HiFi assemblies demonstrated superior performance in key metrics such as NG50 and Longest contig length, where higher values signify better assembly quality. Conversely, HiFi assemblies demonstrated lower values for metrics such as Rduplication, Number of Sequences, Switch, and Hamming errors, indicating superior assembly quality. It’s noteworthy that duplex assemblies had a higher count of T2T contigs and k-mer completeness. The lower NG50 of duplex assemblies may stem from the inherent limitations of handling contained reads within assemblers that employ string graphs, such as Hifiasm (Kamath et al. 2023). The quantitative values for assembly features across replicates are available in the supplementary materials [Figure S8, Table S18].

**Figure 8.**
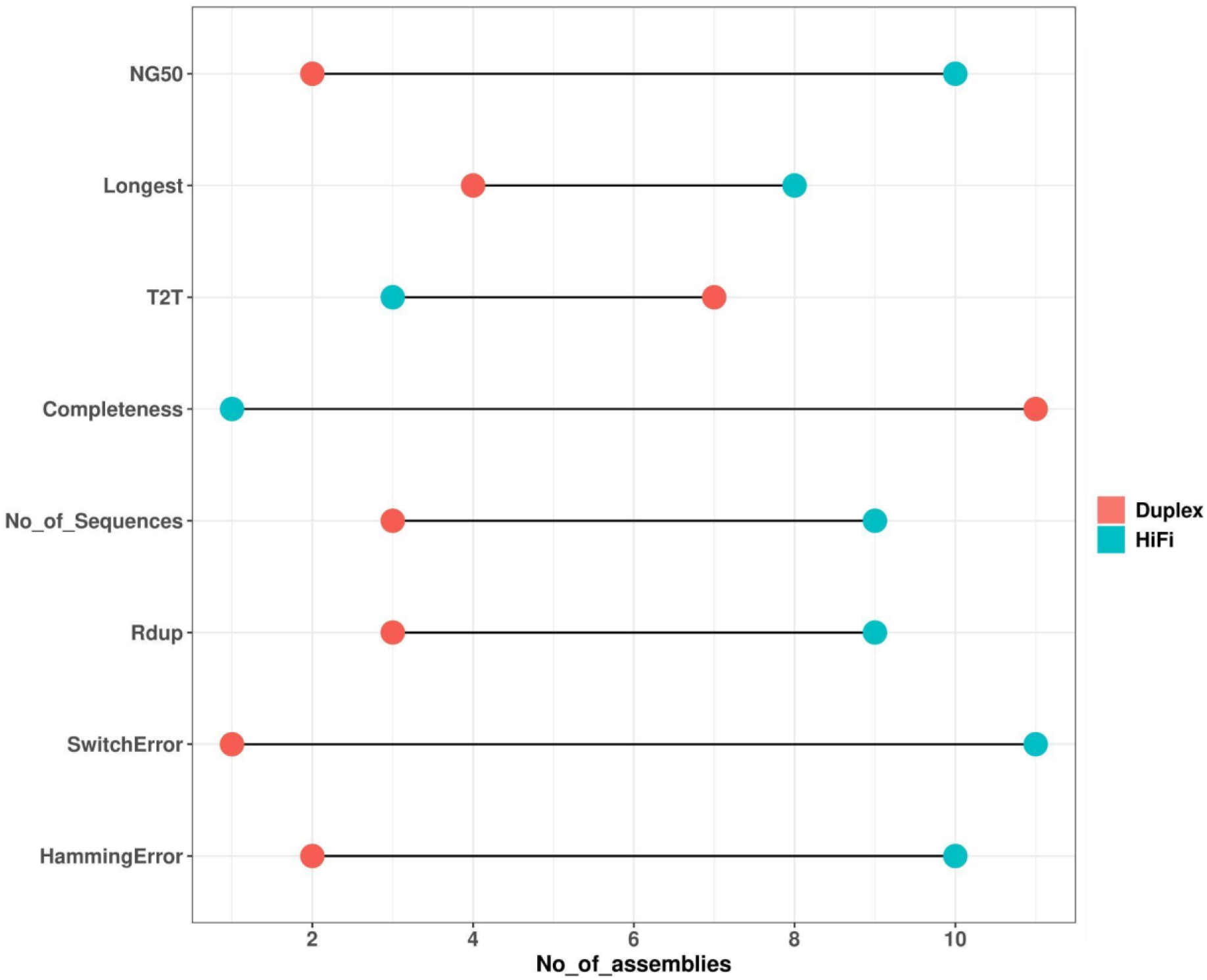
The relative performance of HiFi vs Duplex assemblies from I002C and HG002 using the plateau coverage i.e. 35x HiFi vs 35x Duplex + (30x ULONT and 30x Omni-C). For each assembly feature, we compared the performance of HiFi vs duplex across 12 assemblies (3 replicates * 2 samples [I002C and HG002] * 2 haplotypes) and recorded instances where the HiFi assembly outperformed the duplex assembly and vice versa. For example, 10 out of 12 HiFi assemblies had higher NG50 values compared to duplex assemblies.

## Discussion

The DNA sequencing landscape is continually evolving, with advancements in sequencing technologies offering unprecedented opportunities for genomic research. In this study, we conducted a comprehensive analysis of sequencing data obtained from PacBio HiFi, ONT Duplex, ONT Ultralong (ULONT), and Omni-C data in the context of de novo assembling. We aimed to investigate coverage saturation for different data types and their implications for various aspects of de novo assembly, including phasing, genome completeness, and assembly quality.

Our findings provide valuable insights into the optimal sequencing coverage depth required for de novo assembly in large-scale analyses. Through coverage saturation analysis, we observed a positive correlation between sequencing coverage and assembly performance. Notably, assembly contiguity plateaued when the HQLR-only coverage exceeded 35x. Furthermore, the integration of ULONT data significantly enhanced assembly contiguity, particularly for assembling telomere-to-telomere contigs, underscoring the importance of long-range data in improving assembly contiguity. The assembly contiguity plateaus with ULONT coverage exceeding 30x.

We did not involve parental information in generating haplotype-resolved assemblies. Trio binning using parental data facilitates assembly and increases phasing accuracy compared to long-range data phasing (Koren et al. 2018). However, it requires additional effort in the recruitment process and often parental information is not available. Even with high-quality long reads such as HiFi/Duplex and ULONT with substantial coverage, the assembled genome still can have higher switch and hamming errors. Our study demonstrates the efficacy of incorporating long-range chromatin interaction data like Omni-C to address this issue. By leveraging long-range contact information provided by Omni-C, we observed a notable reduction in globally incorrectly phased variants. However, challenges persist in accurately identifying the parental origin of phased contigs, highlighting the inherent ambiguity in long-range data phasing.

Genome completeness and quality assessments revealed marginal improvements with increased coverage, with assemblies incorporating ULONT data exhibiting higher completeness metrics compared to HQLR-only assemblies. Our analysis emphasizes the importance of considering both single-copy gene analysis and k-mer completeness for a comprehensive assessment of genome quality.

The computational demands associated with genomic analysis are substantial, particularly in population-scale studies. Our study highlights the time and memory requirements associated with the assembly process, emphasizing the need for efficient algorithms and computational resources to handle large datasets effectively.

A comparative analysis between PacBio HiFi and ONT Duplex data revealed the superior performance of HiFi data in de novo assembly.

## Methods

### Long read sequencing (LRS) data generation

#### 1) Pacbio data generation

The high molecular weight (HMW) DNA used for PacBio sequencing was extracted using the GentraPuregene kit (Qiagen; #158043) according to the manufacturer’s instructions. Briefly, 1 x 10^^7^ frozen cell pellets from respective EBV cell lines were used as input for extraction. All vortexing steps were replaced with gentle inversion throughout the process, and 300 µl of Qiagen EB buffer was used for elution. Eluted DNA was incubated at 12°C with gentle shaking over a period of 7 to 10 days. To avoid shearing the high molecular weight DNA, wide bore tips with gentle pipetting were used during handling. DNA was stored at 4°C to prevent freeze and thaw cycle. Quantity and purity of extracted HMW DNA were assessed using triplicate concentration measurements from the top, middle, and bottom sections of sample volume, using Qubit dsDNA BR (Broad-Range) assay (Thermofisher Scientific; Q32853) and NanoDrop 2000 spectrophotometer (ThermoFisher Scientific; ND-2000), according to manufacturer’s instructions.

After DNA extraction, DNA fragment lengths were then measured using TapeStation 4200 (Agilent). Sequencing libraries for each sample were created using the SMRTbell Express Template Prep Kit 2.0 (PacBio) per the manufacturer’s instructions. Libraries were sequenced with one sample per run on a Sequel IIe and Revio System (PacBio). After sequencing, CCS analyses were run using SMRTLink software v10 to produce HiFi reads for each sample.

#### 2) ONT data generation

##### 2.1) High Molecular Weight (HMW) gDNA Extraction (Duplex sequencing)

We obtained 12 x 10^6 frozen cell pellets from established lymphoblastoid cell lines of each trio individual and processed them for HMW gDNA extraction using the Monarch HMW DNA Extraction Kit for Tissue (NEB; T3060). During the extraction, we excluded shaking during all incubation steps to preserve gDNA integrity. Quantity, purity, and integrity of extracted HMW gDNA were assessed using Qubit dsDNA BR assay (Thermofisher Scientific; Q32853), NanoDrop 2000 spectrophotometer (ThermoFisher Scientific; ND-2000), and 15-hour pulsed-field gel electrophoresis runs with the Pippin Pulse system (Sage Science; PPI0200), respectively. Quality-assessed HMW gDNA was then used for ligation-based library preparation with the Ligation Sequencing Kit V14 (Oxford Nanopore Technologies; SQK-LSK114) to generate Duplex sequencing reads.

##### 2.2) Ultra-High Molecular Weight (UHMW) gDNA Extraction (Ultra-long read sequencing)

We processed 15 x 10^6 frozen cell pellets from an established lymphoblastoid cell line for the child sample for UHMW gDNA extraction using the Monarch HMW DNA Extraction Kit for Tissue, following the extraction steps described in the Ultra-Long DNA Sequencing Kit V14 (Oxford Nanopore Technologies; SQK-ULK114) protocol. Quality assessment of UHMW gDNA was performed similarly to HMW gDNA extraction. Quality-assessed UHMW gDNA was then used for transposase-based library preparation and purification with the Ultra-Long DNA Sequencing Kit V14 for the generation of ultra-long sequencing reads.

##### 2.3) Library Preparation and PromethION Sequencing (Duplex and High Duplex)

We sheared 3µg to 7.5µg of extracted HMW gDNA to a target size of 55kb to 60kb and performed size-selective precipitation to remove DNA sizes <25kb. Repaired and end-prepped DNA was then used for library construction with SQK-LSK114 for both duplex and high duplex sequencing approaches. For standard duplex runs, libraries were loaded at 6fmol to 7fmol per load, while for high duplex runs, 7fmol to 55fmol of libraries were loaded and sequenced on PromethION 24 (Oxford Nanopore; PCA100024), R10.4.1 flowcells (Oxford Nanopore) FLO-PRO114M and FLO-PRO114HD respectively.

##### 2.4) Library Preparation and PromethION Sequencing (Ultra-long, UL)

We used 40µg to 45µg of UHMW gDNA for ultra-long read library preparation using SQK-ULK114. Final UL libraries were sequenced on PromethION 24 using FLO-PRO114M flowcells with nuclease flushes performed at 23-hour intervals.

The detailed steps of the entire procedure are outlined in the Supplementary Materials.

#### 3) Omni-C data generation

The Dovetail Omni-C library was prepared using the Dovetail Omni-C™ Proximity Ligation Assay kit (Dovetail Genomics, Scotts Valley, CA, USA), according to the manufacturer’s protocol (version 1.2). Briefly, after sample crosslinking with DSG and formaldehyde, chromatin was digested using a sequence-independent endonuclease and bound to chromatin capture beads. Proximity ligation was performed using a biotin-labeled bridge between the ends of the digested DNA. After reversal crosslinking, the DNA was purified and followed by library preparation. Finally, the biotinylated molecules were captured and amplified before sequencing on the Novaseq 6000 instrument (Illumina, San Diego, CA, USA) with 150 base-length in paired-end mode.

### Data analysis

All commands employed in the analysis are comprehensively listed in the Supplementary Materials file, providing readers with detailed procedures undertaken in this study.

### Reads downsampling

To evaluate coverage saturation for both assembly contiguity and phasing efficiency, we downsampled the reads to various coverages. Reads were randomly subsampled to achieve the desired coverage utilizing Rasusa v0.7.1 (Hall 2022), considering a genome size estimation of 3 gigabases (3g).

### De novo assembly and assessment

Following the insights from the previous benchmark studies (Jarvis et al. 2022; Li and Durbin 2023) and considering the assembler’s capability to produce phased assemblies with constrained data types, we chose Hifiasm v0.19.5-r593 (Cheng et al. 2021; Cheng et al. 2022; Cheng et al. 2023) as the primary assembler. We acknowledge the existence of an alternative, Verkko pipeline (Rautiainen et al. 2023) that can generate telomere-to-telomere phased assemblies by leveraging HiFi/Duplex data and UL data, in tandem with trio or long-range reads. We focused on Hifiasm only due to its shorter running time [Table S1] which we deem could be of high importance in choosing assembly strategy for population genome projects.

To illustrate the significance of diverse data types in producing contiguous and phased assemblies, we executed the assembly process in multiple stages, integrating different data types.

### Assembly Statistics

Assembly contiguity metrics were computed utilizing minigraph v0.20 (Li et al. 2020) and paftools v2.26-r1175 (Li 2018).

### Phasing statistics

The phasing efficiency of an assembly was evaluated in terms of switch error and Hamming error rates with Yak v0.1-r69-dirty (https://github.com/lh3/yak) using parental short reads. Switch error quantifies the frequency of adjacent phased variants incorrectly transitioning between maternal and paternal haplotypes. Meanwhile, the Hamming error rate denotes the total misphased variants within each assembled contig. Phasing statistics were generated for both the haplotypes separately.

### Assembly completeness and quality

To evaluate the impact of coverage variations on the completeness, we employed compleasm v0.2.2 (Huang and Li 2023) to obtain the BUSCO assessment results. Concurrently, we applied a k-mer-based approach for assembly completeness evaluation, employing the KMC tool v3.2.1 (Kokot et al. 2017). Identifying reliable k-mers within the reads followed a previously outlined methodology (Rhie et al. 2020.). The assembly completeness was computed as the fraction of reliable k-mers in the read set that also appeared in the assembly.

## Conclusion

It is essential to recognize the dynamic nature of genomic research and the ongoing evolution of sequencing technologies and analytical methodologies. Through our exploration of various sequencing data types and algorithms, we offer several key insights and recommendations for population-level pan-genome reference generation efforts. We emphasize the pivotal role of integrating high-quality data sources such as Pacbio HiFi/ONT Duplex and ONT ULONT, alongside long-range contract data like Omni-C, to achieve phased telomere-to-telomere level assemblies. In general, HiFi/Duplex coverage of >=20x complemented with 15-20x of ULONT per haplotype and 30x long-range data are essential requisites for attaining high-quality contiguous and phased assembly. We offer our findings as practical guidelines to help users choose sequencing platforms and coverage effectively.

## Data Access

The data utilized in this research are generated as part of an ongoing initiative to develop a telomere-2-telomere diploid assembly of I002C (https://github.com/LHG-GG/I002C / https://github.com/lbcb-sci/I002C). I002C data will be made available to the public upon publication via a cited page. Additionally, in our study, we incorporated publicly available data downloaded from the HPRC (https://humanpangenome.org/data), accessible through AWS.

## Acknowledgments

This study was funded by the UIBR grant from A*STAR, by the European Union through the European Regional Development Fund under the grant KK.01.1.1.01.0009 (DATACROSS), and by the Croatian Science Foundation under the grant IP-2018-01-5886 (SIGMA). The authors thank Oxford Nanopore for providing duplex and ULONT flowcells, as well as PacBio for HiFi flowcells at no cost. We would like to thank Mr Low Hwee Meng and Ms See Ting Leong from GIS Integrated Genomics Platform for their help with generating ONT data. Moreover, we would like to thank Ms Li Zhihui also from GIS Integrated Genomics Platform, for base calling the data as part of ONT-T2T collaboration. Finally, we would like to thank Ms Yao Fei for generating Omni-C data for the I002C T2T project and Ms Sia YY for maintaining the cell line, both from the GIS Laboratory of Human Genomics. This work was supported by the A*STAR Computational Resource Centre through the use of its high performance computing facilities.

## Author Contributions

JJ.L. and M.Š. conceived the project. P.S. designed the pipeline. P.S. and J.L. prepared datasets and conducted the data analysis with the help of F.T. for genome completeness analysis. P.S. wrote the manuscript, and J.L. and F.T. helped with the organization of it. M.Š. and JJ.L. supervised the project and provided mentorship.

## References

Ballouz S, Dobin A, Gillis JA. 2019. Is it time to change the reference genome? Genome Biol 20. 10.1186/s13059-019-1774-4.

Cheng H, Asri M, Lucas J, Koren S, Li H. 2023. Scalable telomere-to-telomere assembly for diploid and polyploid genomes with double graph. arXiv [q-bioGN]. http://arxiv.org/abs/2306.03399 (Accessed February 28, 2024).

Cheng H, Concepcion GT, Feng X, Zhang H, Li H. 2021. Haplotype-resolved de novo assembly using phased assembly graphs with hifiasm. Nat Methods 18: 170–175. 10.1038/s41592-020-01056-5.

Cheng H, Jarvis ED, Fedrigo O, Koepfli K-P, Urban L, Gemmell NJ, Li H. 2022. Haplotype-resolved assembly of diploid genomes without parental data. Nat Biotechnol 40: 1332–1335. 10.1038/s41587-022-01261-x.

Cunningham F, Allen JE, Allen J, Alvarez-Jarreta J, Amode MR, Armean IM, Austine-Orimoloye O, Azov AG, Barnes I, Bennett R, et al. 2022. Ensembl 2022. Nucleic Acids Res 50: D988–D995. 10.1093/nar/gkab1049.

De Coster W, Weissensteiner MH, Sedlazeck FJ. 2021. Towards population-scale long-read sequencing. Nat Rev Genet 22: 572–587. 10.1038/s41576-021-00367-3.

Deng L, Xie B, Wang Y, Zhang X, Xu S. 2022. A protocol for applying a population-specific reference genome assembly to population genetics and medical studies. STAR Protoc 3: 101440. 10.1016/j.xpro.2022.101440.

Du X, Li L, Liang F, Liu S, Zhang W, Sun S, Sun Y, Fan F, Wang L, Liang X, et al. 2022. Robust benchmark structural variant calls of an Asian using state-of-the-art long-read sequencing technologies. Genomics Proteomics Bioinformatics 20: 192–204. 10.1016/j.gpb.2020.10.006.

Ebler J, Ebert P, Clarke WE, Rausch T, Audano PA, Houwaart T, Mao Y, Korbel JO, Eichler EE, Zody MC, et al. 2022. Pangenome-based genome inference allows efficient and accurate genotyping across a wide spectrum of variant classes. Nat Genet 54: 518–525. 10.1038/s41588-022-01043-w.

Eizenga JM, Novak AM, Sibbesen JA, Heumos S, Ghaffaari A, Hickey G, Chang X, Seaman JD, Rounthwaite R, Ebler J, et al. 2020. Pangenome graphs. Annu Rev Genomics Hum Genet 21: 139–162. 10.1146/annurev-genom-120219-080406.

Gao Y, Yang X, Chen H, Tan X, Yang Z, Deng L, Wang B, Kong S, Li S, Cui Y, et al. 2023. A pangenome reference of 36 Chinese populations. Nature 619: 112–121. 10.1038/s41586-023-06173-7.

Garrison E, Sirén J, Novak AM, Hickey G, Eizenga JM, Dawson ET, Jones W, Garg S, Markello C, Lin MF, et al. 2018. Variation graph toolkit improves read mapping by representing genetic variation in the reference. Nat Biotechnol 36: 875–879. 10.1038/nbt.4227.

Hall M. 2022. Rasusa: Randomly subsample sequencing reads to a specified coverage. J Open Source Softw 7: 3941. 10.21105/joss.03941.

Harvey WT, Ebert P, Ebler J, Audano PA, Munson KM, Hoekzema K, Porubsky D, Beck CR, Marschall T, Garimella K, et al. 2023. Whole-genome long-read sequencing downsampling and its effect on variant calling precision and recall. bioRxiv. 10.1101/2023.05.04.539448.

He Y, Chu Y, Guo S, Hu J, Li R, Zheng Y, Ma X, Du Z, Zhao L, Yu W, et al. 2023. T2T-YAO: A telomere-to-telomere assembled diploid reference genome for Han Chinese. Genomics Proteomics Bioinformatics. 10.1016/j.gpb.2023.08.001.

Hickey G, Monlong J, Ebler J, Novak AM, Eizenga JM, Gao Y, Abel HJ, Antonacci-Fulton LL, Asri M, Baid G, et al. 2023. Pangenome graph construction from genome alignments with Minigraph-Cactus. Nat Biotechnol. 10.1038/s41587-023-01793-w.

Huang N, Li H. 2023. compleasm: a faster and more accurate reimplementation of BUSCO. Bioinformatics 39. 10.1093/bioinformatics/btad595.

International Human Genome Sequencing Consortium, Lander ES, Linton LM, Birren B, Nusbaum C, Zody MC, Baldwin J, Devon K, Dewar K, Doyle M, et al. 2001. Initial sequencing and analysis of the human genome. Nature 409: 860–921. 10.1038/35057062.

Jain M, Koren S, Miga KH, Quick J, Rand AC, Sasani TA, Tyson JR, Beggs AD, Dilthey AT, Fiddes IT, et al. 2018. Nanopore sequencing and assembly of a human genome with ultra-long reads. Nat Biotechnol 36: 338–345. 10.1038/nbt.4060.

Jarvis ED, Formenti G, Rhie A, Guarracino A, Yang C, Wood J, Tracey A, Thibaud-Nissen F, Vollger MR, Porubsky D, et al. 2022. Semi-automated assembly of high-quality diploid human reference genomes. Nature 611: 519–531. 10.1038/s41586-022-05325-5.

Kamath SS, Bindra M, Pal D, Jain C. 2023. Telomere-to-telomere assembly by preserving contained reads. bioRxiv. 10.1101/2023.11.07.565066.

Kokot M, Długosz M, Deorowicz S. 2017. KMC 3: counting and manipulating *k*-mer statistics. Bioinformatics 33: 2759–2761. 10.1093/bioinformatics/btx304.

Koren S, Rhie A, Walenz BP, Dilthey AT, Bickhart DM, Kingan SB, Hiendleder S, Williams JL, Smith TPL, Phillippy AM. 2018. De novo assembly of haplotype-resolved genomes with trio binning. Nat Biotechnol 36: 1174–1182. 10.1038/nbt.4277 (Accessed March 4, 2024).

Li H. 2018. Minimap2: pairwise alignment for nucleotide sequences. Bioinformatics 34: 3094–3100. 10.1093/bioinformatics/bty191.

Li H, Durbin R. 2023. Genome assembly in the telomere-to-telomere era. arXiv [q-bioGN]. http://arxiv.org/abs/2308.07877 (Accessed February 28, 2024).

Li H, Feng X, Chu C. 2020. The design and construction of reference pangenome graphs with minigraph. Genome Biol 21. 10.1186/s13059-020-02168-z.

Liao W-W, Asri M, Ebler J, Doerr D, Haukness M, Hickey G, Lu S, Lucas JK, Monlong J, Abel HJ, et al. 2023. A draft human pangenome reference. Nature 617: 312–324. 10.1038/s41586-023-05896-x.

Logsdon GA, Vollger MR, Eichler EE. 2020. Long-read human genome sequencing and its applications. Nat Rev Genet 21: 597–614. 10.1038/s41576-020-0236-x.

Lou H, Gao Y, Xie B, Wang Y, Zhang H, Shi M, Ma S, Zhang X, Liu C, Xu S. 2022. Haplotype-resolved de novo assembly of a Tujia genome suggests the necessity for high-quality population-specific genome references. Cell Syst 13: 321–333.e6. 10.1016/j.cels.2022.01.006.

Ni Y, Liu X, Simeneh ZM, Yang M, Li R. 2023. Benchmarking of Nanopore R10.4 and R9.4.1 flow cells in single-cell whole-genome amplification and whole-genome shotgun sequencing. Comput Struct Biotechnol J 21: 2352–2364. 10.1016/j.csbj.2023.03.038.

Nurk S, Koren S, Rhie A, Rautiainen M, Bzikadze AV, Mikheenko A, Vollger MR, Altemose N, Uralsky L, Gershman A, et al. 2022. The complete sequence of a human genome. Science 376: 44–53. 10.1126/science.abj6987.

Rautiainen M, Nurk S, Walenz BP, Logsdon GA, Porubsky D, Rhie A, Eichler EE, Phillippy AM, Koren S. 2023. Telomere-to-telomere assembly of diploid chromosomes with Verkko. Nat Biotechnol 41: 1474–1482. 10.1038/s41587-023-01662-6.

Rhie A, Nurk S, Cechova M, Hoyt SJ, Taylor DJ, Altemose N, Hook PW, Koren S, Rautiainen M, Alexandrov IA, et al. 2023. The complete sequence of a human Y chromosome. Nature 621: 344–354. https://www.nature.com/articles/s41586-023-06457-y (Accessed February 28, 2024).

Rhie A, Walenz BP, Koren S, Phillippy AM. 2020. Merqury: reference-free quality, completeness, and phasing assessment for genome assemblies. Genome Biol 21. 10.1186/s13059-020-02134-9.

Schneider VA, Graves-Lindsay T, Howe K, Bouk N, Chen H-C, Kitts PA, Murphy TD, Pruitt KD, Thibaud-Nissen F, Albracht D, et al. 2017. Evaluation of GRCh38 and de novo haploid genome assemblies demonstrates the enduring quality of the reference assembly. Genome Res 27: 849–864. https://pubmed.ncbi.nlm.nih.gov/28396521/ (Accessed February 28, 2024).

Sherman RM, Salzberg SL. 2020. Pan-genomics in the human genome era. Nat Rev Genet 21: 243–254. 10.1038/s41576-020-0210-7.

Sur A, Noble WS, Myler PJ. 2022. A benchmark of Hi-C scaffolders using reference genomes and *de novo* assemblies. bioRxiv. 10.1101/2022.04.20.488415.

Vernikos GS. 2020. A review of pangenome tools and recent studies. In The Pangenome, pp. 89–112, Springer International Publishing, Cham.

Wang T, Antonacci-Fulton L, Howe K, Lawson HA, Lucas JK, Phillippy AM, Popejoy AB, Asri M, Carson C, Chaisson MJP, et al. 2022. The Human Pangenome Project: a global resource to map genomic diversity. Nature 604: 437–446. 10.1038/s41586-022-04601-8.

Yang C, Zhou Y, Song Y, Wu D, Zeng Y, Nie L, Liu P, Zhang S, Chen G, Xu J, et al. 2023. The complete and fully-phased diploid genome of a male Han Chinese. Cell Res 33: 745– 761. 10.1038/s41422-023-00849-5.

Yang X, Lee W-P, Ye K, Lee C. 2019. One reference genome is not enough. Genome Biol 20. 10.1186/s13059-019-1717-0.

