## Supplemental Materials for "The Hitchhiker’s Guide to Sequencing Data Types and Volumes for Population-Scale Pangenome Construction"

**Table of Contents**


**Supplemental methods** 2

**ONT data generating** 2

1) High Molecular Weight (HMW) gDNA Extraction (Duplex sequencing) 2

2) Ultra-High Molecular Weight (UHMW) gDNA Extraction (Ultra-long read sequencing) 2

3) Library Preparation and PromethION Sequencing (Duplex and High Duplex) 2

4) Library Preparation and PromethION Sequencing (Ultralong, UL) 3

5) Re-basecalling of Oxford Nanopore Technologies (ONT) pod5 files 3

**ONT base-calling** 4

ONT ULONT 4

Duplex 4

**Reads down-sampling** 4

**De novo assembly** 4

HQLR-only (HiFi/Duplex data) 4

HQLR+ ULONT 4

HQLR+ ULONT + OmniC 4

**Assembly Statistics** 4

**Assembly quality** 5

**Phasing statistics** 5

**Assembly completeness** 5

K-mer base 5

Single-Copy gene analysis 5

**Supplemental figures S1-S7 (attached)** 6

### Supplemental methods

#### ONT data generating

##### 1) High Molecular Weight (HMW) gDNA Extraction (Duplex sequencing)

12 x 10ˆ6 frozen cell pellets from established lymphoblastoid cell line obtained as input to HMW gDNA extraction with the Monarch HMW DNA Extraction Kit for Tissue (NEB; T3060), according to the manufacturer's protocol, with exclusion of shaking during all incubation steps to allow for maximum gDNA integrity. Quantity and purity of extracted HMW gDNA were assessed using triplicate measurements from the top, middle and bottom sections of sample volume, using Qubit dsDNA BR (Broad-Range) assay (Thermofisher Scientific; Q32853) and NanoDrop 2000 spectrophotometer (ThermoFisher Scientific; ND-2000), according to manufacturer's instructions. gDNA integrity was assessed via 15-hour pulsed-field gel electrophoresis runs with the Pippin Pulse system (Sage Science; PPI0200), using the 5 - 430kb pre-set protocol on 0.75% Lonza SeaKem agarose gels casted at 12 x 14cm, with the midi gel box (Sage Science; PGB1000) and 0.5X KBB buffer (Sage Science; KBB1001). Quality-assessed HMW gDNA was carried forward to ligation-based library preparation using the Ligation Sequencing Kit V14 (Oxford Nanopore Technologies; SQK-LSK114) for the generation of Duplex sequencing reads.

##### 2) Ultra-High Molecular Weight (UHMW) gDNA Extraction (Ultra-long read sequencing)

15 x 10ˆ6 frozen cell pellets from established lymphoblastoid cell line for child sample were obtained as input to UHMW gDNA extraction with the Monarch HMW DNA Extraction Kit for Tissue, following extraction steps described in the Ultra-Long DNA Sequencing Kit V14 (Oxford Nanopore Technologies; SQK-ULK114) protocol. Final UHMW gDNA elution was performed with ultra-long read workflow specific ONT Extraction Elution Buffer (EEB), to a final volume of 760µl. gDNA quality assessment was performed as per HMW gDNA for Duplex sequencing described in the previous section. Quality-assessed UHMW gDNA was carried forward to transposase-based library preparation and purification using the Ultra-Long DNA Sequencing Kit V14, according to the manufacturer's protocol, for the generation of ultra-long sequencing reads.

##### 3) Library Preparation and PromethION Sequencing (Duplex and High Duplex)

3µg to 7.5µg of extracted HMW gDNA was sheared to a target size of 55kb to 60kb using the Megaruptor 3 Shearing Kit (Diagenode; E07010003) on the Megaruptor® 3 instrument (Diagenode; B06010003), according to recommended manufacturer's settings. Sheared gDNA size distribution was assessed using the Genomic DNA ScreenTape (Agilent Technologies; 5067-5366, 5067-5365) using 4200 TapeStation System (Agilent Technologies; G2991BA). Size-selective precipitation was performed on sheared gDNA for progressive depletion of DNA sizes <25kb, with either a Short Read Eliminator (SRE) kit (PacBio; SKU 102-208-300) or Short Fragment Eliminator (SFE) Expansion kit (Oxford Nanopore Technologies; EXP-SFE001), according to respective manufacturer's instructions. Size distribution of size-selected gDNA was assessed on the Genomic DNA ScreenTape before proceeding to library construction with SQK-LSK114 for both Duplex and high Duplex sequencing approaches.

1.8µg to 4.7µg of size-selected gDNA input were subjected to a combined DNA repair and end-prep reaction using FFPE Repair Mix and Ultra II End repair/dA-tailing Module (New England Biolabs, M6630 and E7546), with incubation for 20 minutes at 20°C and 10 minutes at 65°C. Repaired and end-prepped DNA was purified using 0.4X AMPure XP beads, with elution in nuclease-free water, followed by Ligation Adapter (LA) ligation incubation for 30 minutes at room temperature, increased from the default of 10 minutes. Purification of the adapted library was performed using 0.4X AMPure XP beads, two Long Fragment Buffer (LFB) washes, and elution in ONT Elution Buffer (EB). All DNA elution steps described were performed with incubation at 37°C for 10 minutes to aid recovery of HMW DNA libraries. Final libraries were quantified on both Qubit Fluorometer and Genomic DNA ScreenTape on the 4200 TapeStation System to assess concentration and size distribution, respectively.

Libraries were sequenced on PromethION 24 (Oxford Nanopore; PCA100024), R10.4.1 flowcells (Oxford Nanopore; FLO-PRO114M and FLO-PRO114HD), for standard Duplex and high Duplex runs respectively, each employing different library loading molarity strategies.

For standard Duplex, libraries were loaded at 6fmol to 7fmol per load, with nuclease flushes performed at 20 to 23-hour intervals, using the flowcell wash kit (Oxford Nanopore Technologies; EXP-WSH004), according to manufacturer's instructions. Two to three nuclease flushes were performed for the sequencing runs, for a total of three to four library loadings per flowcell, over sequencing duration between 72 to 100 hours, depending on monitored flowcell performance. For high Duplex runs, 7fmol to 55fmol of libraries were loaded, while maintaining the nuclease flush intervals and sequencing durations as described above.

##### 4) Library Preparation and PromethION Sequencing (Ultralong, UL)

40µg to 45µg of UHMW gDNA reconstituted in 750µl of EEB was used as input to three times scaled Nanopore ultra-long read library preparation using SQK-ULK114. For each library prepared, 6µl ONT Fragmentation Mix (FRA) was diluted with 244µl FRA Dilution Buffer (FDB) and added to 750µl UHMW DNA, followed in quick succession with gentle pipetting with P1000 wide bore tips to ensure even distribution. The DNA tagmentation reaction was incubated at room temperature for 10 minutes, followed by inactivation at 75°C, for 10 minutes, and cooling on ice for 10 minutes. Rapid sequencing adaptor attachment was conducted with the addition of 5µl of Rapid Adapter (RA) to the cooled tagmentation reaction, followed by gentle pipetting with P1000 wide bore tips and incubation at room temperature for 30 minutes. Library purification was performed using the Precipitation Star (PS) and Precipitation buffer (PTB) approach as per the manufacturer's protocol, with overnight elution of the purified UL library. Final UL libraries were assessed for yield using the Qubit dsDNA BR assay and divided into three aliquots of approximately 90µl each, for PromethION sequencing as per SQK-ULK114 protocol.

UL libraries were sequenced on PromethION 24 using FLO-PRO114M flowcells, with pore scan intervals set at default of 1.5 hours. Nuclease flushes were performed according to the manufacturer's instructions, using the flowcell wash kit, EXP-WSH004, at 23-hour intervals, followed by reloading of the UL library aliquot, for a total of two flushes and three loads per library.

##### 5) Re-basecalling of Oxford Nanopore Technologies (ONT) pod5 files

The output pod5 files from MiniKNOW are re-basecalled with dorado v0.3.0 and converted to bam file with samtools v1.16.1. Flowcells which come from Duplex protocol had the bam file further processed with duplex_tools v0.3.2 to identify the Duplex reads. The table below shows the exact command for the list of protocols used. The detailed commands used for basecalling are provided in Supplementary Materials.

#### ONT base-calling

##### ONT ULONT

```
dorado basecaller -r --emit-sam --modified-bases 5mCG_5hmCG \
--min-qscore 7 dna_r10.4.1_e8.2_400bps_sup@v4.1.0 pod5_folder/ | \
samtools view --bam -Sh -O BAM -o ultra-long.unmapped_reads.bam -
```

##### Duplex

1.
```
dorado basecaller --emit-moves -r --emit-sam \
--modified-bases 5mCG_5hmCG --min-qscore 7 \
dna_r10.4.1_e8.2_400bps_hac@v4.2.0 pod5_folder/ | \
samtools view --bam -Sh -O BAM -o duplex.unmapped_reads.bam -
```

2.
```
duplex_tools pair --output_dir pairs_from_bam/ duplex.unmapped_reads.bam
```

3.
```
duplex_tools split_pairs duplex.unmapped_reads.bam \
pod5_folder/pod5s_splitduplex/
```

4.
```
cat pod5s_splitduplex/*_pair_ids.txt > split_duplex_pair_ids.txt
```

5.
```
dorado duplex --emit-fastq --pairs pairs_from_bam/pair_ids_filtered.txt \
--min-qscore 7 dna_r10.4.1_e8.2_400bps_sup@v4.2.0 pod5_folder/ \
> duplex_orig.fastq
```

6.
```
dorado duplex --emit-fastq --pairs split_duplex_pair_ids.txt \
--min-qscore 7 dna_r10.4.1_e8.2_400bps_sup@v4.1.0 pod5s_splitduplex/ \
>  duplex_splitduplex.fastq
```

#### Reads down-sampling

```
rasusa -o {output.fq} -i {input.fq} -c {desired_coverage} -g 3000000000
```

#### De novo assembly

##### HQLR-only (HiFi/Duplex data)

```
hifiasm -t {threads} -o {output} {input.fq}
```

##### HQLR+ ULONT

```
hifiasm -t {threads} -o {output} --ul {ulont.fq} {input.fq}
```

##### HQLR+ ULONT + OmniC

```
hifiasm -t {threads} -o {output} --ul {ulont.fq} --h1 {read1.fq} --h2 \
{read2.fq} {input.fq}
```

#### Assembly Statistics

```
minigraph -x asm –show-unmap=yes -t {threads} -K1.9g {reference} \
```

```
{input.fa}  >{output.paf}

paftools.js asmstat {reference.fa.fai} {output.paf} >{out.assembly.stats}
```

#### Assembly quality

Creating the k-mer database:

```
yak count -k21 -t {threads}-b37 -o {output.yak} \
 <(zcat {input.fq.gz}) <(zcat {input.fq.gz})
```

Calculating the assembly QV

```
yak qv -t {threads} {output.yak} {assembly.fa} >{assembly.fa_qv}
```

#### Phasing statistics

```
yak trioeval -t {threads} {paternal.yak} {maternal.yak)\
 {haplotype.fa} >{haplotype_phasing.txt}
```

#### Assembly completeness

##### K-mer base

```
kmc -k21 -m100 -t {threads} -fm -ci1 {assembly.fa} {assembly_kmers} $PWD

kmc -k21 -m100 -t {threads} -fq -ci1 {reads.fq} {reads_kmers} $PWD

kmc_tools transform {reads_kmers} histogram {reads_histogram.txt}

kmc_tools simple {reads_kmers} -ci{realiable_kmer_count} \
{assembly_kmers} {read_only_reliable_kmers}
```

##### % assembly completeness:

```
1- { read_only_reliable_kmers / all_reads_reliable_kmers} * 100
```

##### Single-Copy gene analysis

```
compleasm run -a {input.fa} -o {output} -t {threads} -l {mammalia_lineage}\
 -L {library_path} -m {Busco_mode}
```

### Supplemental figures S1-S7 (attached)

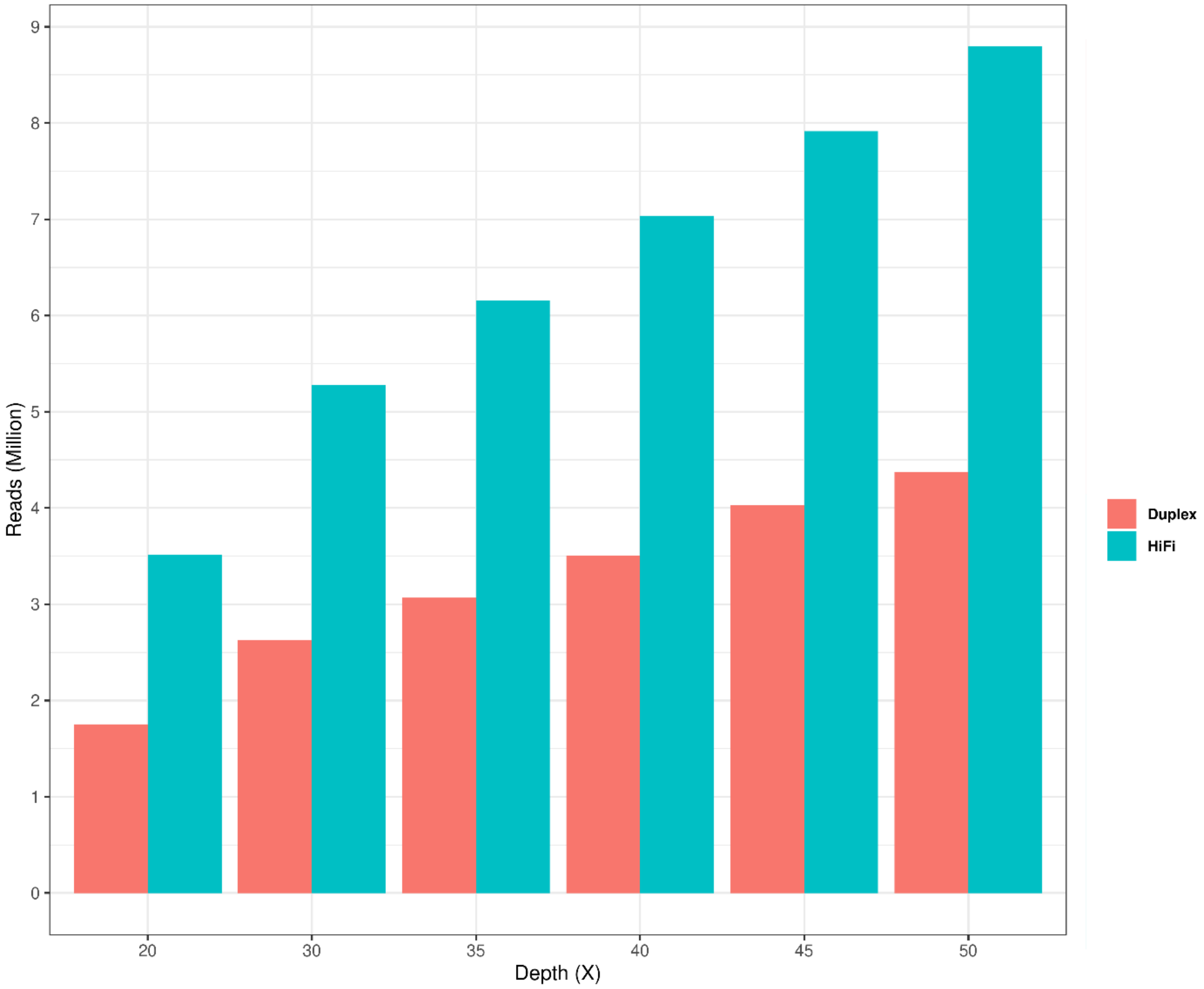


**Figure S1:** Comparison of reads required for a given depth: PacBio HiFi vs ONT Duplex.

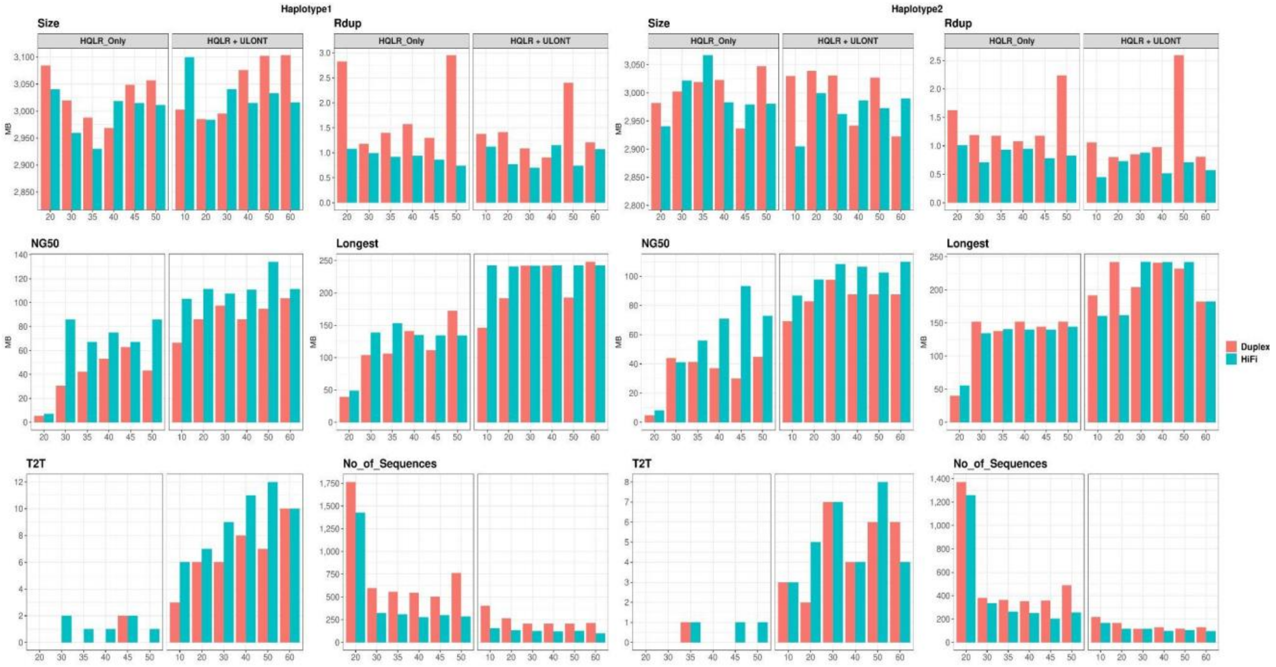


**Figure S2:** Haplotype-wise comparison of assembly performance vs data coverage. Assembly contiguity features like NG50, T2T contigs, and Longest contigs tend to increase with coverages while reaching a plateau at 35x for HQLR-Only and 30x for ULONT data. Similarly, “No_of_Sequences” decreases with an increase in coverage reaching a plateau around the same coverage compared to contiguity features.

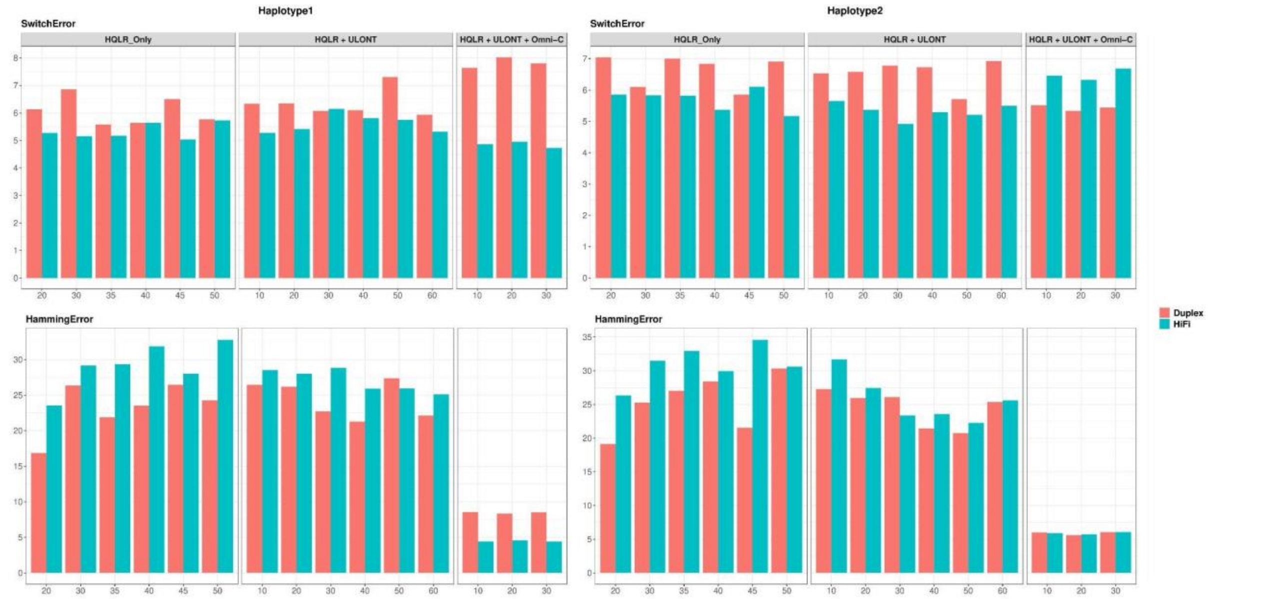


**Figure S3:** Haplotype-wise comparison of switch error and hamming error. Global phasing efficiency measured in terms of hamming error significantly improved with the introduction of long-range data.

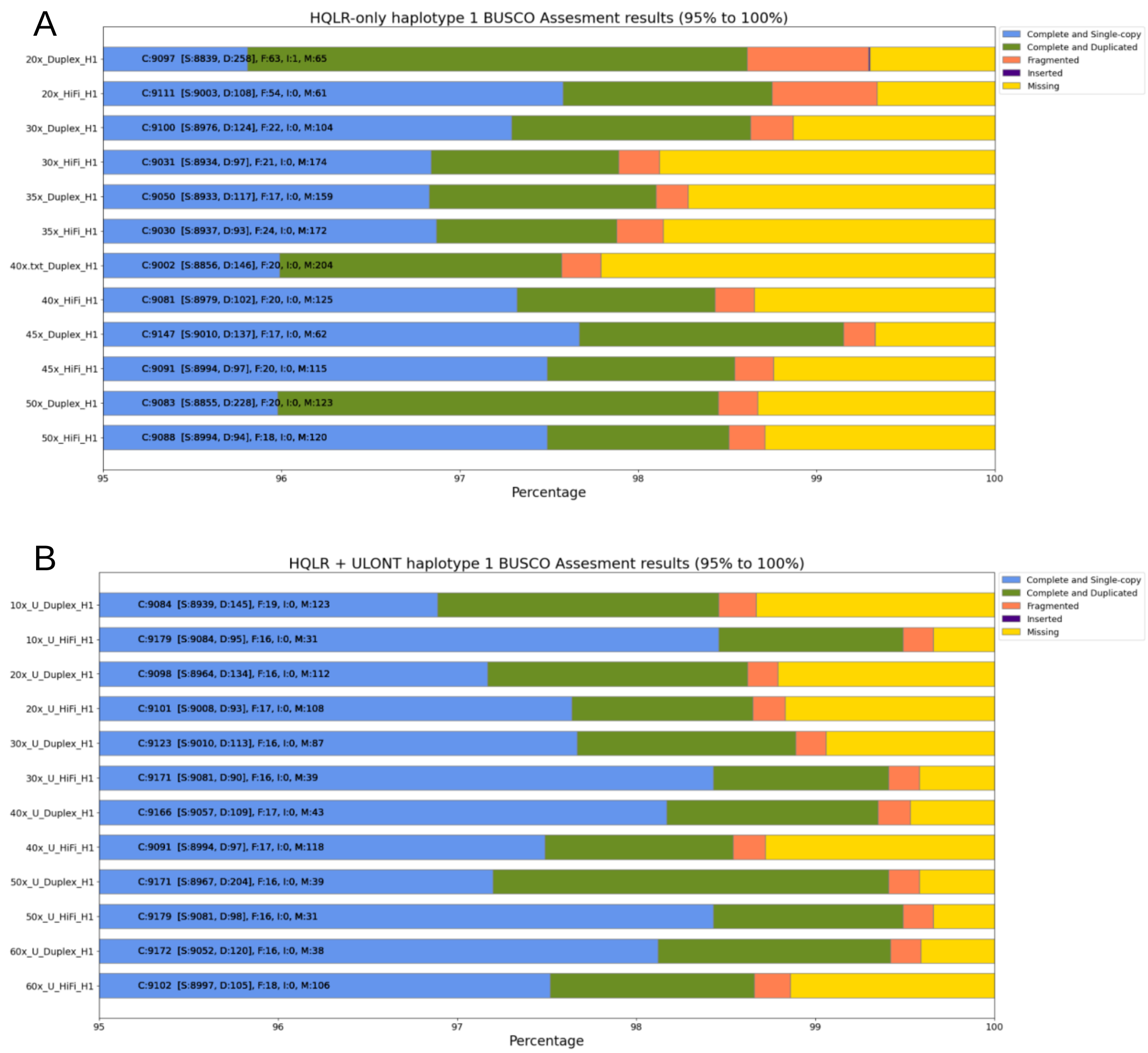


**Figure S4:** Comparison of single copy gene analysis of A) HQLR_only vs B) HQLR + ULONT for haplotype1.

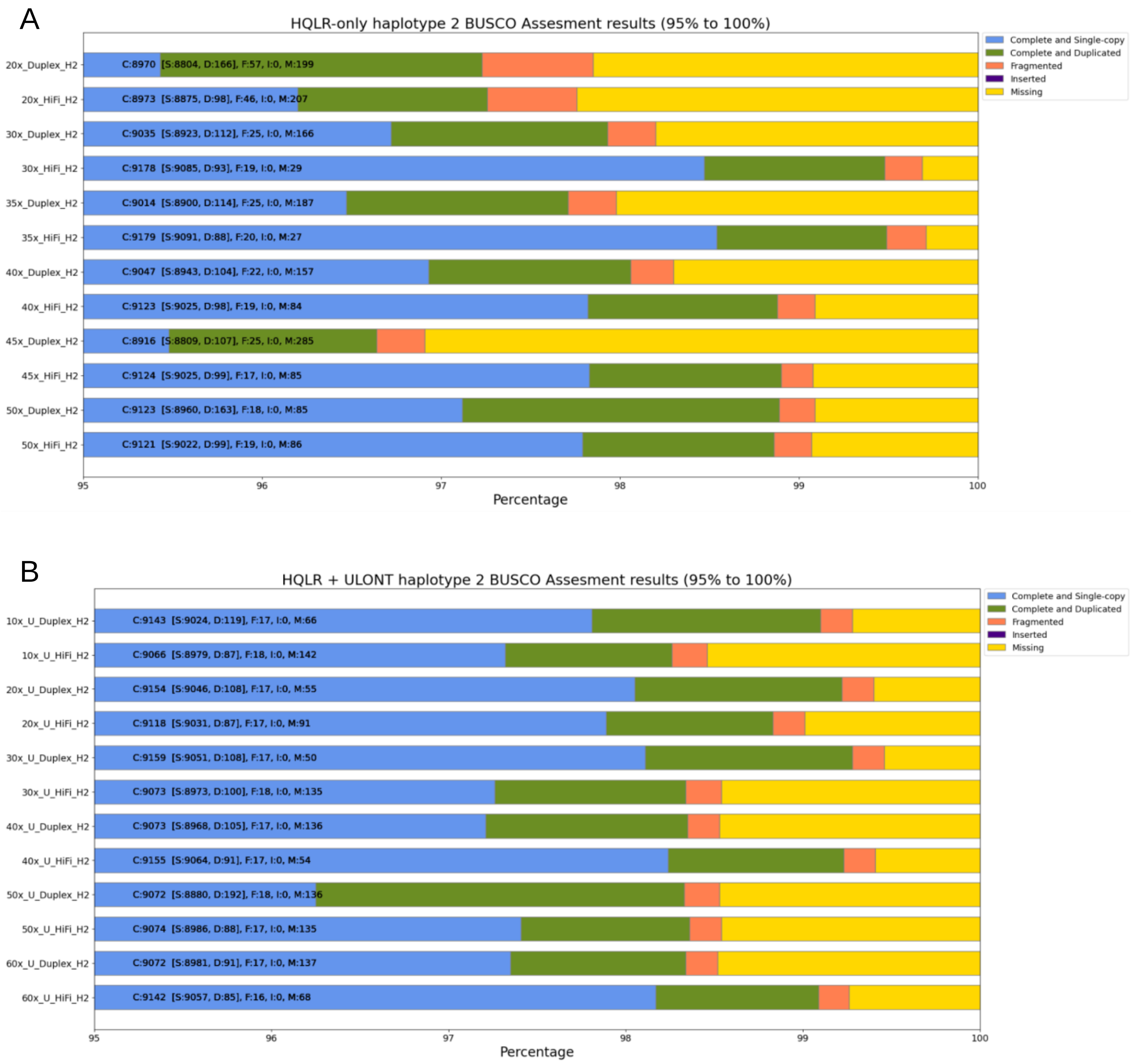


**Figure S5:** Comparison of single copy gene analysis of A) HQLR_only vs B) HQLR + ULONT for haplotype2.

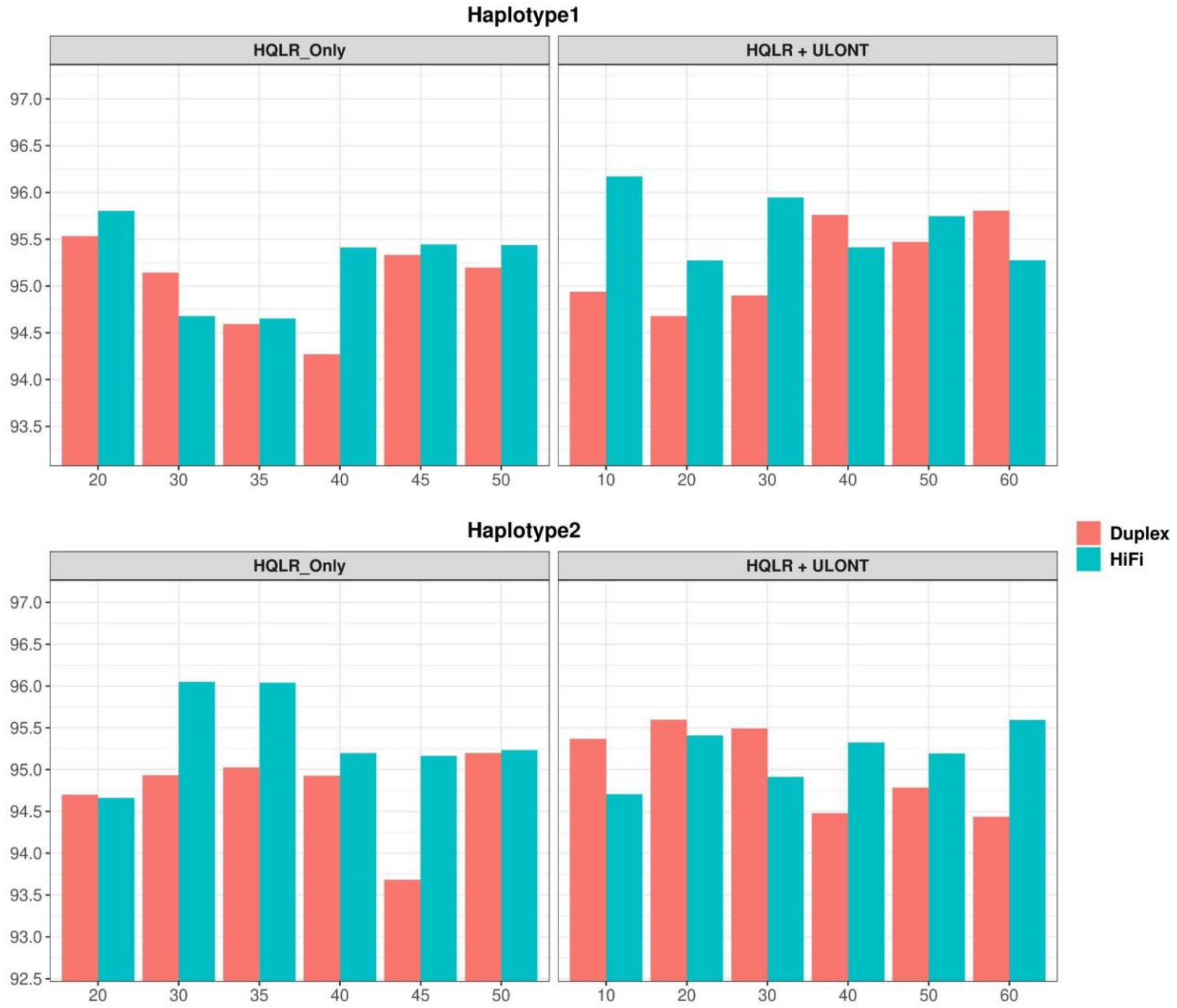


**Figure S6:** Haplotype-wise comparison of k-mer-based assembly completeness. Completeness is not significantly influenced by the sequencing depth while the average completeness of HQLR_ULONT assemblies is marginally higher than HQLR_only assemblies.

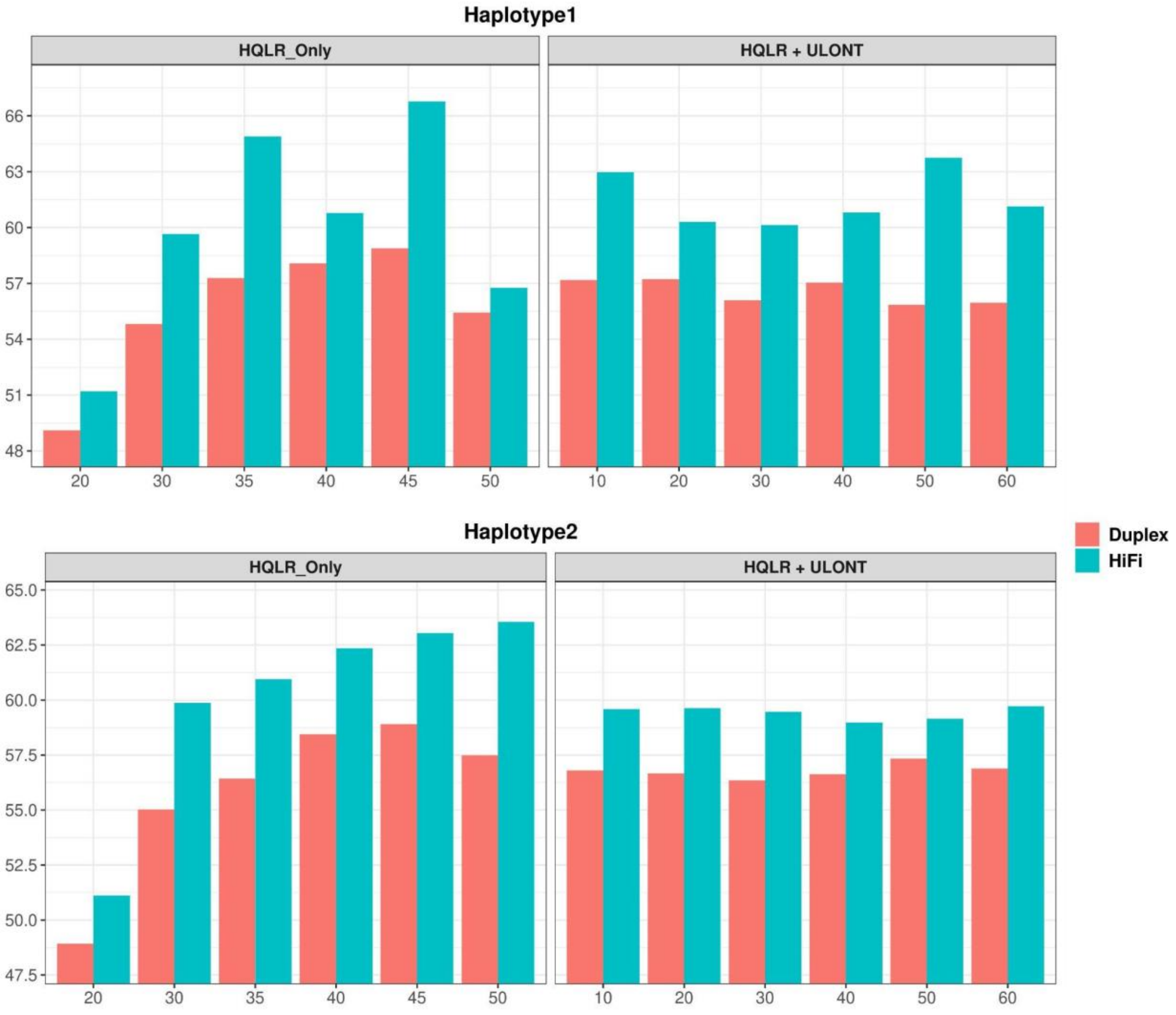


**Figure S7:** Haplotype-wise comparison of assembly QV. Assembly quality of HQLR_only assemblies showed a positive correlation with data coverage.

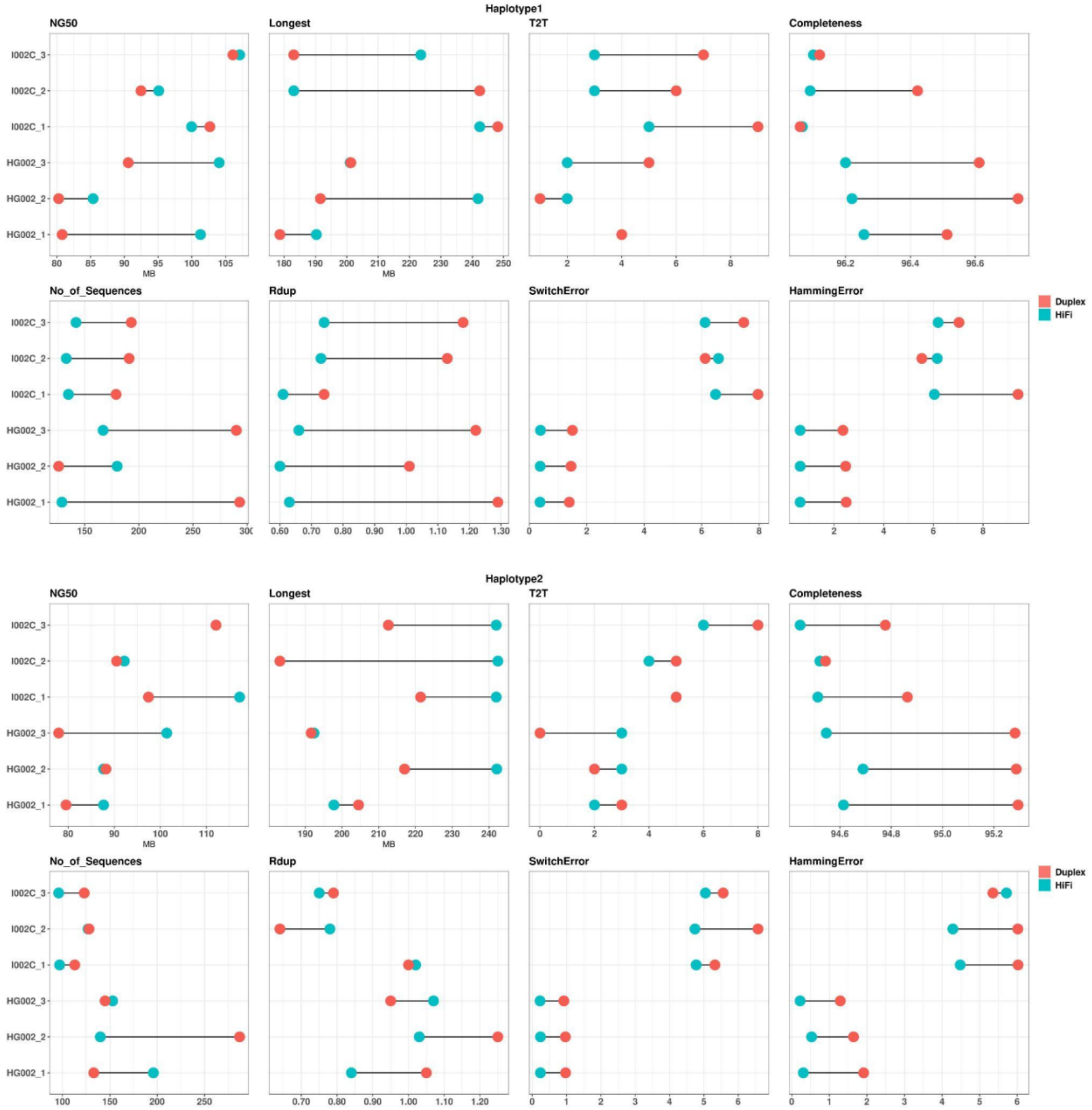


**Figure S8:** Haplotype-wise comparison of assembly features between HiFi vs Duplex dataset. Higher values of assembly features like NG50, Longest, T2T and completeness, coupled with lower value of No_of_Sequences, Rdup, Switch and Hamming error, collectively indicate a better quality assembly.
